# Synergistic Control of Mitochondrial Dynamics and Function by ERAD and Autophagy in Brown Adipocytes

**DOI:** 10.1101/2025.06.14.659625

**Authors:** Xinxin Chen, Siwen Wang, Mauricio Torres, Sijie Hao, Shengyi Sun, Ling Qi

## Abstract

Mitochondrial quality control is essential for maintaining cellular energy homeostasis, particularly in brown adipocytes where dynamic mitochondrial remodeling supports thermogenesis. Although the SEL1L-HRD1 endoplasmic reticulum (ER)-associated degradation (ERAD) pathway and autophagy are two major proteostatic systems, how these pathways intersect to regulate mitochondrial integrity in metabolically active tissues remains poorly understood. Here, using adipocyte-specific genetic mouse models combined with high-resolution 2D and 3D ultrastructural imaging technologies, we reveal an unexpected synergy between SEL1L-HRD1 ERAD and autophagy in maintaining mitochondrial structure and function in brown adipocytes. Loss of ERAD alone triggers compensatory autophagy, whereas combined deletion of both pathways (double knockout, DKO) results in severe mitochondrial abnormalities, including the accumulation of hyperfused megamitochondria penetrated by ER tubules, even under basal room temperature conditions. These phenotypes are absent in mice lacking either pathway individually or in SEL1L-IRE1α DKO, highlighting the pathway-specific coordination between ERAD and autophagy. Mechanistically, dual loss of ERAD and autophagy induces ER expansion, excessive ER-mitochondria contact, upregulation of mitochondria-associated membrane (MAM) tethering proteins, impaired calcium transfer, and defective mitochondrial turnover. As a result, DKO adipocytes accumulate dysfunctional mitochondria, exhibit respiratory deficits, and fail to sustain thermogenesis. Collectively, our study uncovers a cooperative and previously unrecognized mechanism of mitochondrial surveillance, emphasizing the critical role of ERAD-autophagy crosstalk in preserving mitochondrial integrity and thermogenic capacity in brown fat.

**One-sentence summary:** Our study uncovers a previously unrecognized synergy between SEL1L-HRD1 ERAD and autophagy that is essential for preserving mitochondrial integrity and thermogenic capacity in brown adipocytes, revealing new opportunities for targeting mitochondrial dysfunction in metabolic disease.

## INTRODUCTION

Mitochondria are central hubs of cellular metabolism, playing essential roles in energy production, thermogenesis, and cell survival. Their functional integrity is maintained through continuous remodeling by tightly regulated processes including fusion, fission, biogenesis, and selective degradation, collectively termed ‘mitochondrial quality control’ ^1,2^. This surveillance is especially critical in brown adipose tissue (BAT), a highly metabolically active organ specialized in non-shivering thermogenesis ^3,4^. Brown adipocytes are densely populated with mitochondria, whose morphology and activity dynamically respond to environmental cues such as temperature and nutrient status ^5^. Disruption of mitochondrial quality control in BAT impairs thermogenic capacity and contributes to metabolic dysfunction, obesity, and related disorders ^3,6,7^; however, the underlying molecular mechanism remains vague.

Cellular proteostasis and organelle quality control are maintained by two highly conserved, principal degradative systems: autophagy and the endoplasmic reticulum (ER)-associated degradation (ERAD) pathway. While autophagy has been extensively studied ^8–11^, ERAD remains relatively underappreciated, despite its critical role in health and disease ^12–14^. Autophagy is a conserved lysosomal degradation pathway essential for maintaining cellular homeostasis by removing damaged organelles (including the ER and mitochondria), misfolded protein aggregates, and other cytoplasmic components ^8–11^. Macroautophagy, the most extensively studied form, involves the formation of double-membrane autophagosomes that enclose cytoplasmic cargo and subsequently fuse with lysosomes for degradation ^15^. Autophagy plays both constitutive and stress-responsive roles, being upregulated in response to nutrient starvation, hypoxia, and other cellular stresses ^16^. Selective forms of autophagy, such as ER-phagy and mitophagy, enable organelle-specific turnover to preserve organellar function and metabolic homeostasis ^17–20^. Dysregulated autophagy has been implicated in diverse human pathologies, including neurodegenerative diseases, cancer, and metabolic syndromes ^21^.

In BAT, autophagy plays complex and context-dependent roles. Brown adipocyte-specific knockout (KO) models of core autophagy genes have revealed critical and context-dependent roles for autophagy in regulating mitochondrial homeostasis, thermogenesis, and systemic energy metabolism. Deletion of *Atg7* ^22,23^, *Atg5* or *Atg12* ^24^ using Cre drivers like aP2-Cre, Ucp1-Cre or Ucp1-CreERT2 impairs mitophagy and alters mitochondrial turnover, increased mitochondrial content and enhanced thermogenic potential. In contrast, inducible deletion of *Atg3* or *Atg16L1* in mature adipocytes using Adipoq-CreERT2 results in mitochondrial dysfunction, oxidative stress, and insulin resistance ^25^. These findings suggest that while autophagy restricts excessive mitochondrial accumulation during adipogenesis, it is indispensable for mitochondrial quality and metabolic health in mature BAT.

Unlike autophagy, ERAD targets misfolded proteins in the ER for proteasomal degradation in the cytosol ^26–30^. The SEL1L-HRD1 complex constitutes the most conserved branch of ERAD ^31–34^. In this complex, SEL1L serves as an essential adaptor for the E3 ubiquitin ligase HRD1, where SEL1L acts as a scaffold, linking substrates to HRD1 and stabilizing HRD1 ^35–39^. Deletion of *Sel1L* or *Hrd1* in mice results in embryonic lethality or premature death ^38,40–42^, and loss of ERAD function in specific cell types leads to functional impairment and disease in mice ^12–14,43–49^. In humans, mutations in *SEL1L* or *HRD1* cause ERAD-associated neurodevelopmental disorder with onset in infancy (ENDI) syndrome ^50,51^. We previously identified SEL1L-HRD1 ERAD as a key regulator of mitochondrial dynamics in brown adipocytes during cold exposure, in part by modulating ER-mitochondria contacts or mitochondria-associated membranes (MAMs) ^52^. MAMs are specialized ER-mitochondria contact sites that coordinate mitochondrial dynamics, lipid and calcium exchange, and autophagosome biogenesis ^53–60^. ERAD deficiency leads to increased levels of the MAM-resident protein Sigma non-opioid intracellular receptor 1 (SIGMAR1), mitochondrial hyperfusion and the accumulation of pleomorphic mitochondria with impaired respiratory function following acute cold stress ^52^. While both ERAD and autophagy contribute to ER and mitochondrial homeostasis ^14^, how they intersect to maintain mitochondrial integrity *in vivo* remains unclear.

Since damaged mitochondria are typically cleared by mitophagy ^61^, and that ERAD deficiency activates autophagy in various cell types including white adipocytes and pancreatic β cells ^62,63^, we hypothesized that ERAD and autophagy may act cooperatively to preserve mitochondrial quality in BAT. Using adipocyte-specific KO mouse models combined with high-resolution 2D and 3D ultrastructural imaging technologies, we demonstrate a synergistic interplay between SEL1L-HRD1 ERAD and autophagy in regulating mitochondrial morphology, function, and calcium fluxes in brown adipocytes. This coordination occurs through modulation of MAMs and ehanced clearance of damaged mitochondria. Notably, this cooperative regulation is absent in brown adipocytes deficient in both ERAD and IRE1α, a key sensor of the unfolded protein response ^12,64^. Our findings uncover a previously unappreciated, integrated role for ERAD and autophagy in mitochondrial quality control and thermogenic homeostasis *in vivo*.

## RESULTS

### Activation of autophagy in *Sel1L*-deficient brown adipocytes

Our previous work demonstrated that loss of *Sel1L* activates autophagy in pancreatic β cells and white adipocytes ^62,63^. To assess whether a similar response occurs in brown adipocytes, we analyzed brown adipose tissues (BAT) from brown adipocyte-specific *Sel1L*-deficient (*Sel1L^Ucp1Cre^*) mice. Compared to WT littermates, *Sel1L^Ucp1Cre^* mice exhibited a twofold increase in LC3-II levels, the ATG7-dependent, lipidated form of LC3 associated with autophagosome membrane (**Figure 1A**). Transmission electron microscopy (TEM) revealed a greater number of autophagic-like structures (yellow arrows), often enclosing mitochondria-like organelles (asterisks), in *Sel1L^Ucp1Cre^* BAT (**Figure S1A**). Moreover, we observed a marked increase in mitochondrial ubiquitination in purified mitochondria from *Sel1L^Ucp1Cre^* BAT (**Figure 1B**), a hallmark of elevated mitophagy. These findings establish that *Sel1L* deficiency triggers autophagy in brown adipocytes, consistent with previous findings in other metabolically active cell types.

**Figure 1.**
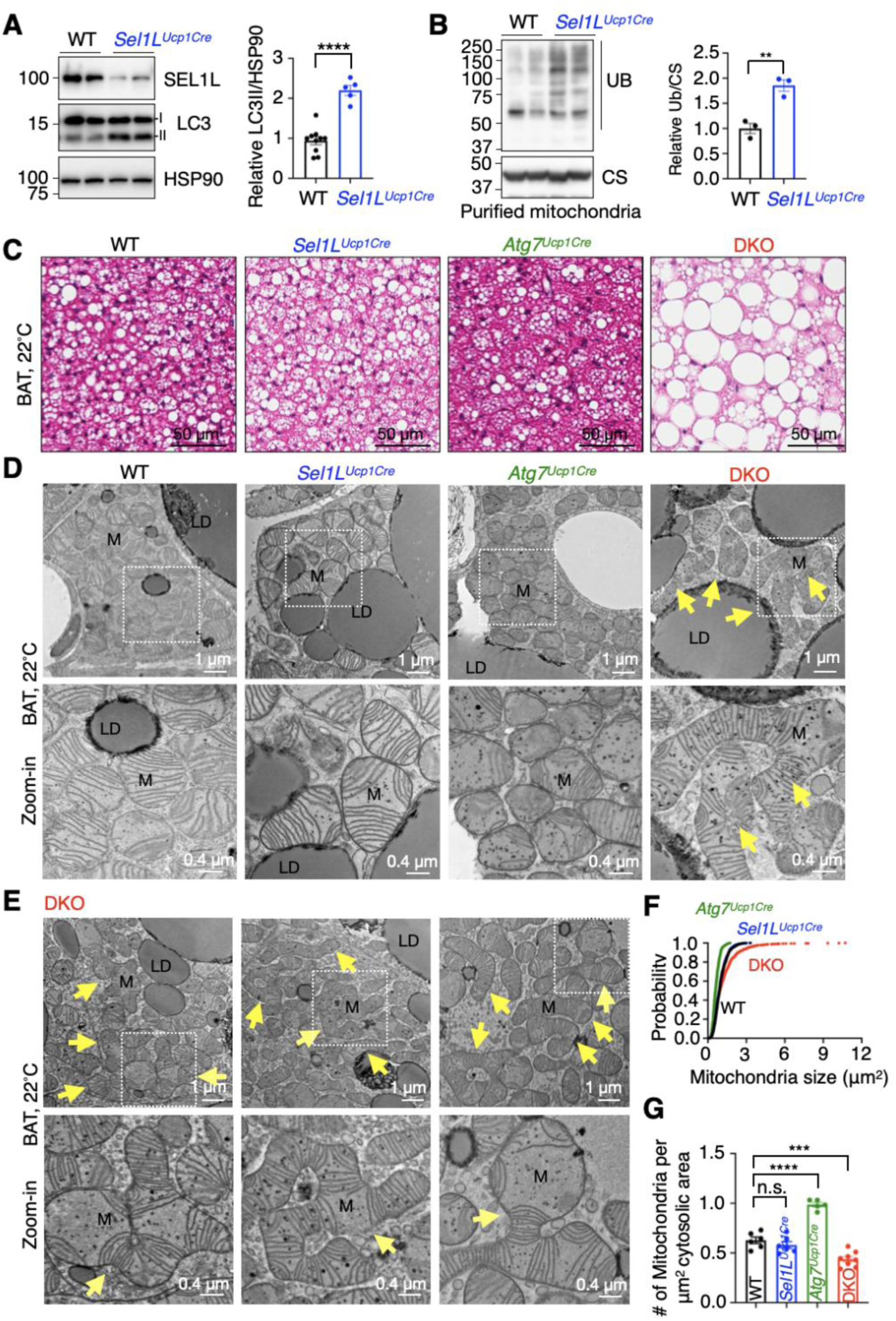
The synergistic role of SEL1L-HRD1 ERAD and autophagy in regulating mitochondrial morphology in brown adipocytes under basal conditions. (A) Immunoblot analysis of LC3 in BAT from 12-week-old WT *and Sel1L^Ucp1Cre^* littermates, with quantification of relative LC3-II normalized to HSP90 shown on the right. n = 10 for WT and 5 for *Sel1L^Ucp1Cre^* mice, Student’s *t*-test. (B) Immunoblot analysis of ubiquitin (UB) in purified mitochondria from BAT of 12-week-old WT *and Sel1L^Ucp1Cre^* littermates, with quantification of relative UB normalized to citrate synthase (CS) shown on the right. n = 3 per group, Student’s *t-* test. (C) Representative hematoxylin and eosin (H&E) staining of BAT from 12-week-old male littermates housed at room temperature (RT). n = 3 mice per group. (D-E) Representative TEM images of BAT from 12-week-old male littermates housed at RT. Arrows point to megamitochondria. LD, lipid droplet; M, mitochondrion. n = 3-5 mice per group. (F) Quantification of cumulative probability distribution of mitochondrial size. n = 874 mitochondria for WT, 747 for *Sel1L^Ucp1Cre^*, 556 for *Atg7^Ucp1Cre^*, and 1189 for DKO, 3-5 mice per group. (G) Quantification of mitochondrial density (number of mitochondria per μm^2^ cytosolic area). n = 5-8 fields, 2-3 mice per group. Data are mean ± SEM. n.s., not significant; **, *p* < 0.01; ***, *p* < 0.001; ****, *p* < 0.0001.

### Generation of mice with combined ERAD and autophagy deficiency in brown adipocytes

To investigate the physiological significance of autophagy in the context of *Sel1L* deficiency, we generated brown adipocyte-specific *Sel1L* and *Atg7* double-deficient (*DKO*) mice (*Sel1L ^f/f^;Atg7^f/f^;Ucp1-Cre*) by crossing *Sel1L^f/f^;Atg7^f/f^* mice with *Ucp1-Cre* transgenic mice. Age-matched WT, *Sel1L^Ucp1Cre^*, and *Atg7^Ucp1Cre^* mice were included as controls. While single KO mice grew comparably to WT littermates, DKO mice of both sexes exhibited significant growth retardation (**Figure S1B**). The etiology of this phenotype remains unclear but contrasts with our previous observations in *Sel1L;Atg7^AdipoqCre^ (Sel1L ^f/f^;Atg7^f/f^;Adipoq-Cre)* mice, in which deletion of the same two genes in all adipocytes (under the control of an adipocyte-specific *Adiponectin-Cre)* did not result in growth abnormalities ^63^. These results suggest that the growth defect in DKO mice may be due to leaky expression of *Ucp1-Cre* in other tissues such as kidneys, adrenal glands, thymus, and brain, as recently reported ^65^.

At room temperature, BAT weights were significantly increased in both *Atg7^Ucp1Cre^* and DKO mice compared to those of WT and *Sel1L^Ucp1Cre^* mice, regardless of normalization to body weight (**Figure S1C-D**). While single deletions of *Sel1L* or *Atg7* did not alter brown adipocyte morphology, combined deficiency led to pronounced lipid droplet hypertrophy, as revealed by hematoxylin and eosin (H&E) staining and Perilipin 1 immunostaining (**Figure 1C and S1E-F**). These changes were consistent across both sexes (data not shown).

Western blot analysis confirmed efficient deletion of SEL1L and ATG7 in DKO BAT and demonstrated loss of autophagic activity, as indicated by p62 accumulation and reduced LC3-II formation (**Figure S2A**). Consistent with IRE1α being a known ERAD substrate ^66^, IRE1α protein levels were markedly elevated ∼10-fold in *Sel1L^Ucp1Cre^* and 44-fold in *DKO* BAT compared to *WT* (**Figure S2B**). Splicing of *Xbp1* mRNA, a downstream target of IRE1α, was modestly increased, with spliced *Xbp1* comprising 21% of total *Xbp1* transcripts in *DKO* BAT vs. ∼ 2-4% in single KO BAT (**Figure S2B**). Similarly, the UPR sensor PERK were elevated by ∼3-fold in *Sel1L^Ucp1Cre^* and ∼6-fold in DKO BAT, accompanied by increased phosphorylation of its downstream target eIF2α (**Figure S2C**). Despite this moderate ER stress response, we did not observe evidence of apoptosis, as cleaved caspase-3 remained undetectable in DKO BAT (**Figure S2D**).

### Megamitochondria formation in DKO brown adipocytes at room temperature

Building on our prior work demonstrating a role for SEL1L-HRD1 ERAD in regulating mitochondrial morphology during cold exposure in brown adipocytes ^52^, we investigated how concurrent loss of SEL1L-HRD1 ERAD and autophagy affects mitochondrial architecture under basal (room temperature) conditions. Using transmission electron microscopy (TEM), we observed no significant differences in mitochondrial morphology or density between WT and *Sel1L^Ucp1Cre^* BAT, consistent with previous findings (**Figure 1D**, quantified in **Figure 1F-G**). In contrast, *Atg7 ^Ucp1Cre^* BAT exhibited smaller, more densely packed mitochondria relative to WT controls. Strikingly, brown adipocytes from DKO mice displayed dramatic mitochondrial enlargement and reduced mitochondrial density (arrows, **Figure 1D-E**, quantified in **Figure 1F-G**). Similar megamitochondrial structures were also observed in BAT from *Adiponectin-Cre-* driven DKO mice (arrows, **Figure S1G**), suggesting that this phenotype is a robust consequence of dual ERAD-autophagy loss.

### 3D FIB-SEM reveals mitochondrial hyperfusion in DKO BAT

These ultrastructural findings were corroborated by high-resolution confocal imaging of the mitochondrial outer membrane protein TOMM20 and matrix protein PDH, which further confirmed the presence of enlarged mitochondria in DKO BAT (white arrows, **Figure 2A and S3A,** quantified in **Figure S3B**). To further investigate mitochondrial architecture, we performed focused ion beam-scanning electron microscopy (FIB-SEM). We used convolutional neural network–based image segmentation to reconstruct individual mitochondria at ∼5 nm isotropic resolution in all dimensions (x, y, z), as previously described ^52^. Three-dimensional reconstruction revealed that mitochondria in *Sel1L^Ucp1Cre^* and *Atg7 ^Ucp1Cre^* BAT remained largely spherical and morphologically similar to those in WT BAT (**Figure 2B; Video S1-S3**). In stark contrast, mitochondria in DKO BAT were significantly enlarged and highly pleomorphic, displaying extensive fusion events (**Figure 2B; Video S4,** quantified in **Figure S3C**). Notably, we observed a substantial population of hyperfused megamitochondria in DKO BAT, with the largest reaching a volume of ∼35 μm³— approximately 15-fold greater than the average WT mitochondrial volume of 2.3 μm³. Pseudo-color segmentation of individual mitochondria revealed that this single megamitochondrion alone accounted for ∼25% of the total mitochondrial volume within the imaged DKO field (**Figure 2B; Video S4**). These findings demonstrate that combined loss of ERAD and autophagy drives aberrant mitochondrial fusion and the accumulation of pleomorphic megamitochondria in brown adipocytes, even in the absence of external stressors such as cold exposure.

**Figure 2.**
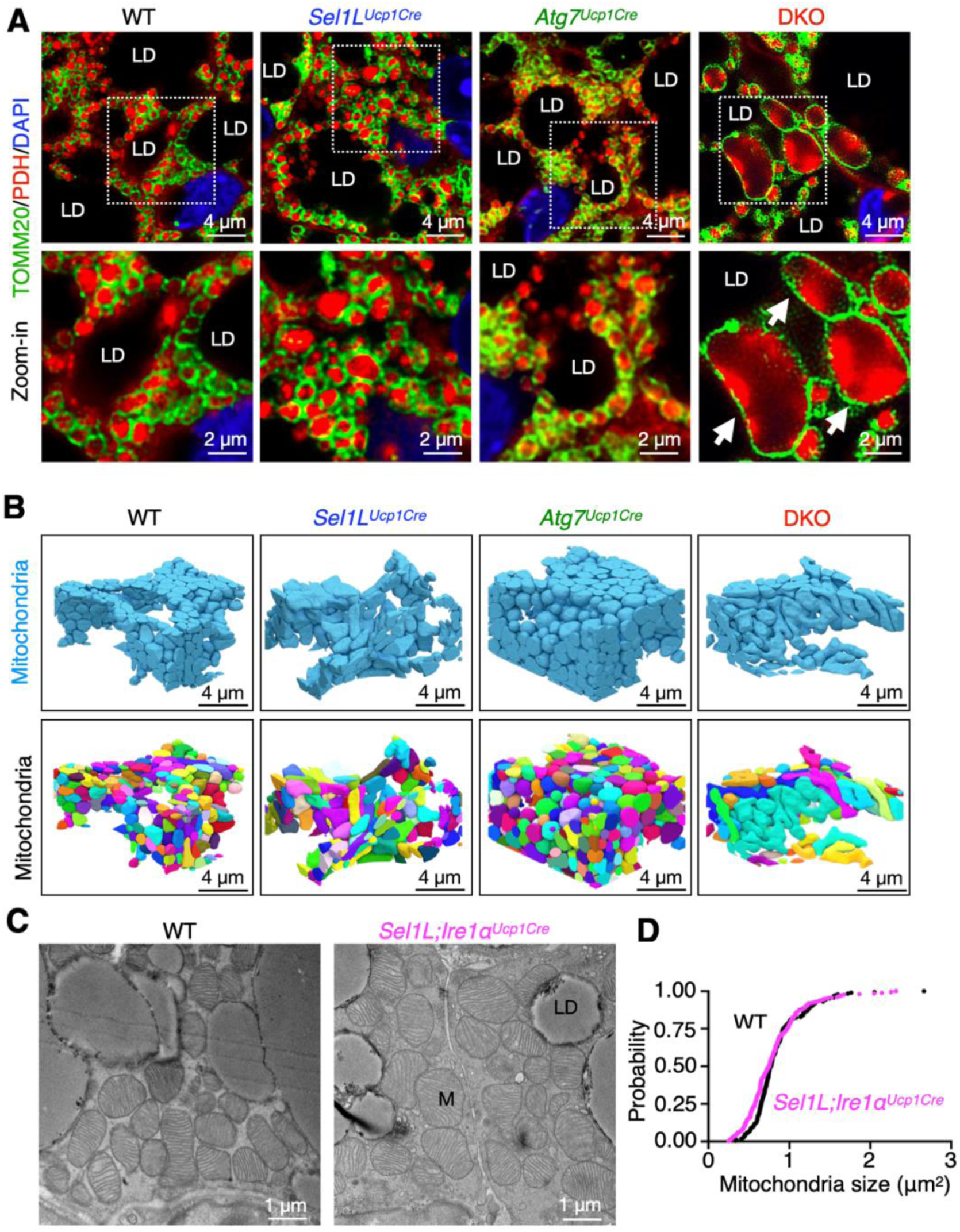
Formation of megamitochondria in *Sel1L;Atg7*-deficient brown adipocytes, but not in *Sel1L;Ire1a*-deficient brown adipocytes. (A) Representative confocal immunofluorescent images of TOMM20 (mitochondrial outer membrane protein) and pyruvate dehydrogenase (PDH, mitochondrial matrix protein) staining in BAT sections from 12-week-old male littermates. Arrows point to megamitochondria. LD, lipid droplet. n = 3 mice per group. (B) Representative FIB-SEM and 3D tomography images of mitochondria in BAT from 12-week-old male littermates. n = 1096 slices for WT, 1425 for *Sel1L^Ucp1Cre^*, 1290 for *Atg7^Ucp1Cre^*, and 1039 for DKO, 5 nm/slice. (C) Representative TEM images of BAT from 10-week-old WT and *Sel1L;Ire1α^Ucp1Cre^* male littermates. LD, lipid droplet; M, mitochondrion. n = 2 mice per group. (D) Quantification of cumulative probability distribution of mitochondrial size. n = 255 and 187 mitochondria for WT and *Sel1L;Ire1α^Ucp1Cre^* mice. n = 2 mice per group

### Combined *Sel1L*-*Ire1a* deficiency does not promote megamitochondria formation

To determine whether the mitochondrial phenotype observed in *Sel1L-Atg7* deficiency is specific to the loss of these two pathways, we generated brown adipocyte-specific *Sel1L* and *Ire1a* DKO (*Sel1L^Ucp1Cre^;Ire1a^Ucp1Cre^*) mice along with their WT littermates (**Figure S3D**). This model was particularly informative given the marked elevation of IRE1α protein levels in *Sel1L^Ucp1Cre^* BAT (**Figure S2B and S3D**). As expected, deletion of IRE1α abolished *Xbp1* mRNA splicing in both *Ire1a^Ucp1Cre^* and *Sel1L^Ucp1Cre^;Ire1a^Ucp1Cre^* BAT (**Figure S3D**). Importantly, *Sel1L^Ucp1Cre^;Ire1a^Ucp1Cre^* mice exhibited normal growth compared to WT littermates (**Figure S3E**) and did not display any evidence of megamitochondria in BAT at 10 weeks of age under room temperature conditions (**Figure 2C**, quantified in **Figure S3G,** and **Fig. S3F**). These data suggest that mitochondrial remodeling in *Sel1L-Atg7* DKO BAT is independent of IRE1α and is unlikely to be a downstream consequence of ER stress.

### Formation of megamitochondria with perforating ER tubules in DKO brown adipocytes

In DKO brown adipocytes, we observed a striking ultrastructural phenotype: in addition to pleomorphic megamitochondria, many of these mitochondria were traversed by perforating tubules (red arrows, **Figure 3A** and **S4A**). These remarkable tubular structures did not puncture the mitochondrial membranes; rather, they exhibited a distinct tri-laminar membrane organization, with two layers derived from the mitochondria (white arrows, **Figure 3A**). Notably, mitochondrial cristae radiated outward from the contact sites of these structures (**Figure 3A** and **S4A**), consistent with a proposed role for the ER in lipid transfer to mitochondria ^55,67^. Occasionally, we observed a unique configuration in which a donut-shaped megamitochondrion encircled a smaller spherical mitochondrion (asterisk), with intervening perforating tubules (red arrows, Figure 3B), suggesting a complex interplay between mitochondrial dynamics and ER contacts.

**Figure 3.**
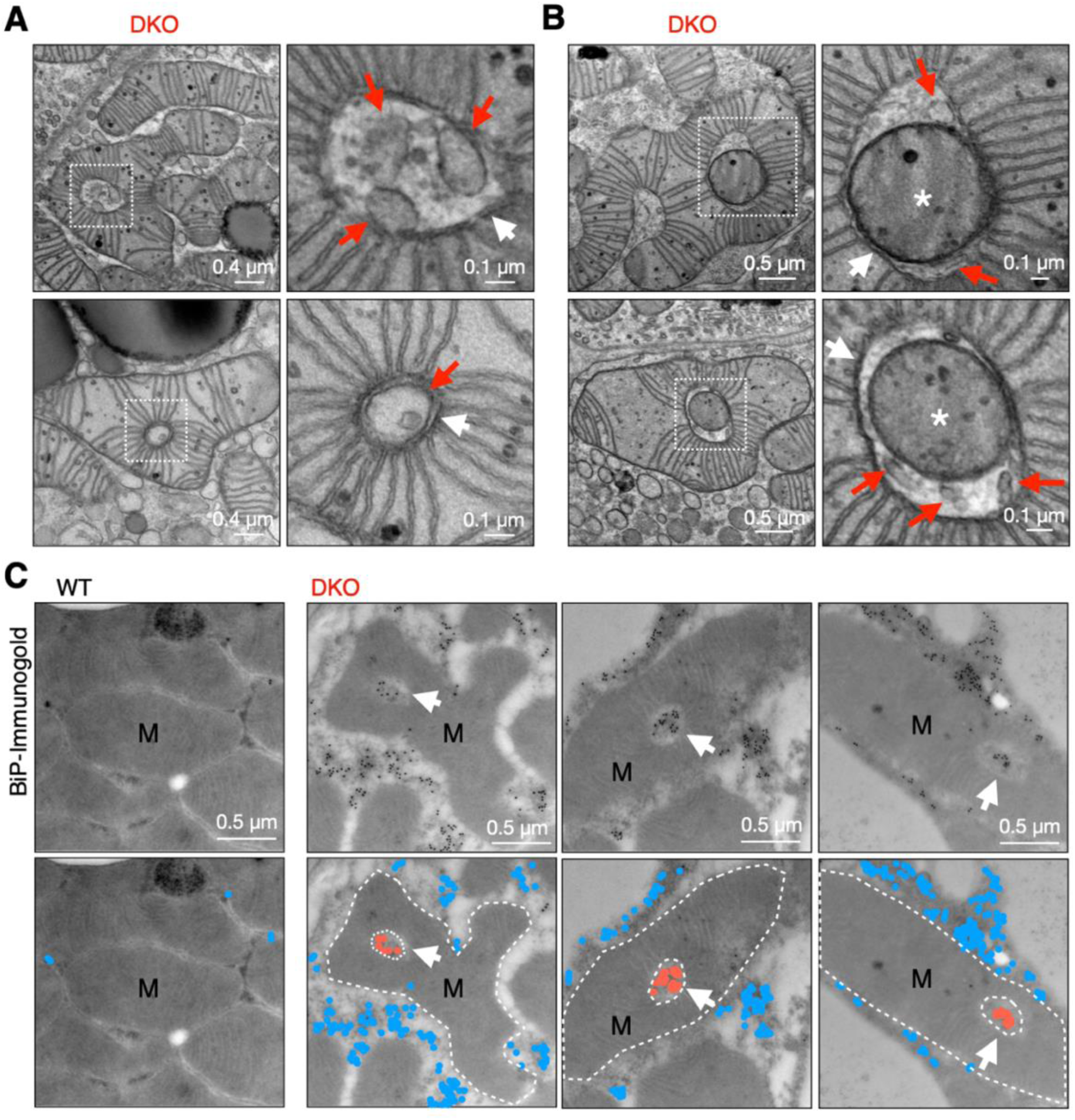
ERAD and autophagy deficiency leads to the formation of megamitochondria with perforating ER tubules and mitochondria under RT. (A) Representative TEM images of BAT from 12-week-old DKO mice showing megamitochondria wrapping around the tubular structures. Red arrows point to mitochondrion-perforating tubules. White arrows point to mitochondrial membranes. n = 5 mice. (B) Representative TEM images of BAT from 12-week-old DKO mice showing megamitochondrion wrapping around the smaller mitochondrion mediated by tubular structures. Red arrows point to mitochondrion-perforating tubules. White arrows point to mitochondrial outer membrane. Asterisk, smaller mitochondrion. n = 5 mice. (C) Representative TEM images of BAT from 12-week-old DKO male mice following BiP-specific immunogold labelling, with pseudo colored blue and red dots shown below. White dotted lines outline megamitochondria. Arrows point to mitochondrion-perforating ER tubules. M, mitochondrion. n = 2 mice per group.

To confirm the identity of these tubules, we performed immunogold labeling for the ER chaperone BiP, which revealed dense clusters of BiP-positive signals surrounding and penetrating the mitochondria (white arrows, **Figure 3C and S4B**), verifying their ER origin. These findings demonstrate that *Sel1L*-*Atg7* deficiency promotes the formation of pleomorphic megamitochondria penetrated by ER-derived tubules in brown adipocytes under room temperature conditions—a phenotype distinct from that seen in *Sel1L^Ucp1Cre^* BAT, where similar structures only emerge under acute cold exposure ^52^. Together, these results highlight a synergistic role of SEL1L-HRD1 ERAD and autophagy in regulating mitochondrial architecture, mitochondrial quality control, and ER–mitochondria interactions.

### ERAD and autophagy synergistically control 3D ER architecture and volume

To elucidate the mechanisms underlying the dramatic mitochondrial alterations in DKO brown adipocytes, we next examined ER architecture, as both SEL1L-HRD1 ERAD and autophagy are known to influence ER morphology and homeostasis ^19,62,68^. TEM revealed modest ER dilation in *Sel1L^Ucp1Cre^* and increased ER sheet formation in *Atg7^Ucp1Cre^* brown adipocytes **(Figure 4A)**. In contrast, DKO adipocytes displayed extensive expansion of rounded tubular ER structures **(Figure 4A)**, confirmed as the ER via BiP immunogold labeling (**Figure 4B and S5A**). Three-dimensional FIB-SEM further revealed that ER occupied ∼6.3% of the cytoplasmic volume in WT adipocytes (excluding lipid droplets, nuclei, and blood vessels) (**Figure 4C, S5B; Video S1**). *Sel1L* or *Atg7* deficiency alone led to a moderate increase in ER volume, whereas combined deletion in DKO adipocytes resulted in a pronounced expansion to ∼14% of the cytosolic volume (**Figure 4C; Video S2-4**), along with increased ER luminal space (arrows, **Figure 4D**). These findings demonstrate a synergistic role for ERAD and autophagy in maintaining ER architecture and controlling ER network volume in brown adipocytes.

**Figure 4.**
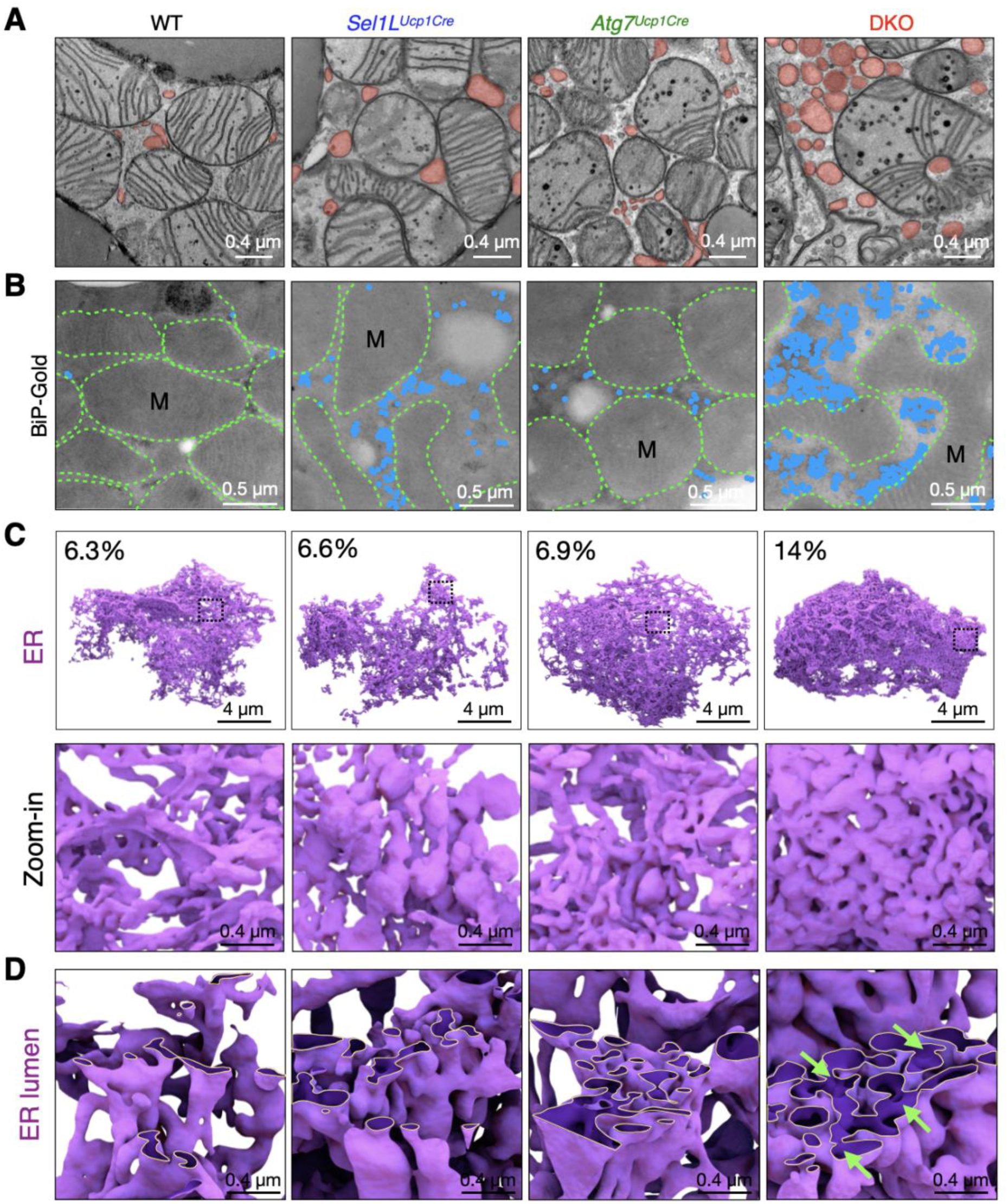
ERAD and autophagy synergistically control the ER architecture in brown adipocytes. (A) Representative TEM images of BAT showing the ER (red) from 12-week-old male littermates. n = 3-5 mice per group. (B) Representative TEM images of BAT following BiP-specific immunogold labelling with pseudo-colored blue dots from 12-week-old male littermates. M, mitochondrion. Green dotted lines outline mitochondrial membranes. n = 2 mice per genotype. (C) 3D reconstruction of FIB-SEM images of BAT showing ER from 12-week-old male littermates, with quantification of the ER volume as the percentage of the total cytosolic volume, excluding the volumes of nucleus and lipid droplets in each group. (D) Magnified view of the ER lumen. Yellow lines outline ER membrane. Green arrows indicate the enlarged ER lumen.

### Enhanced ER-mitochondria contacts in DKO brown adipocytes

Given the critical role of MAMs in mitochondrial dynamics and function ^54,55^, we next explored how SEL1L-HRD1 ERAD and autophagy influence MAM architecture in BAT from mice housed at room temperature. In WT BAT, MAMs appeared as thin ER sheets along mitochondria, with an average distance of 5-25 nm (yellow arrows, **Figure 5A**). A modest increase in MAM number was observed in both *Sel1L^Ucp1Cre^* and *Atg7^Ucp1Cre^* BAT (yellow arrows, **Figure 5A**, and quantified in **Figure 5B**), whereas DKO BAT exhibited a striking accumulation of MAMs per mitochondrion (yellow arrows, **Figure 5A**, quantified in **Figure 5B**).

**Figure 5.**
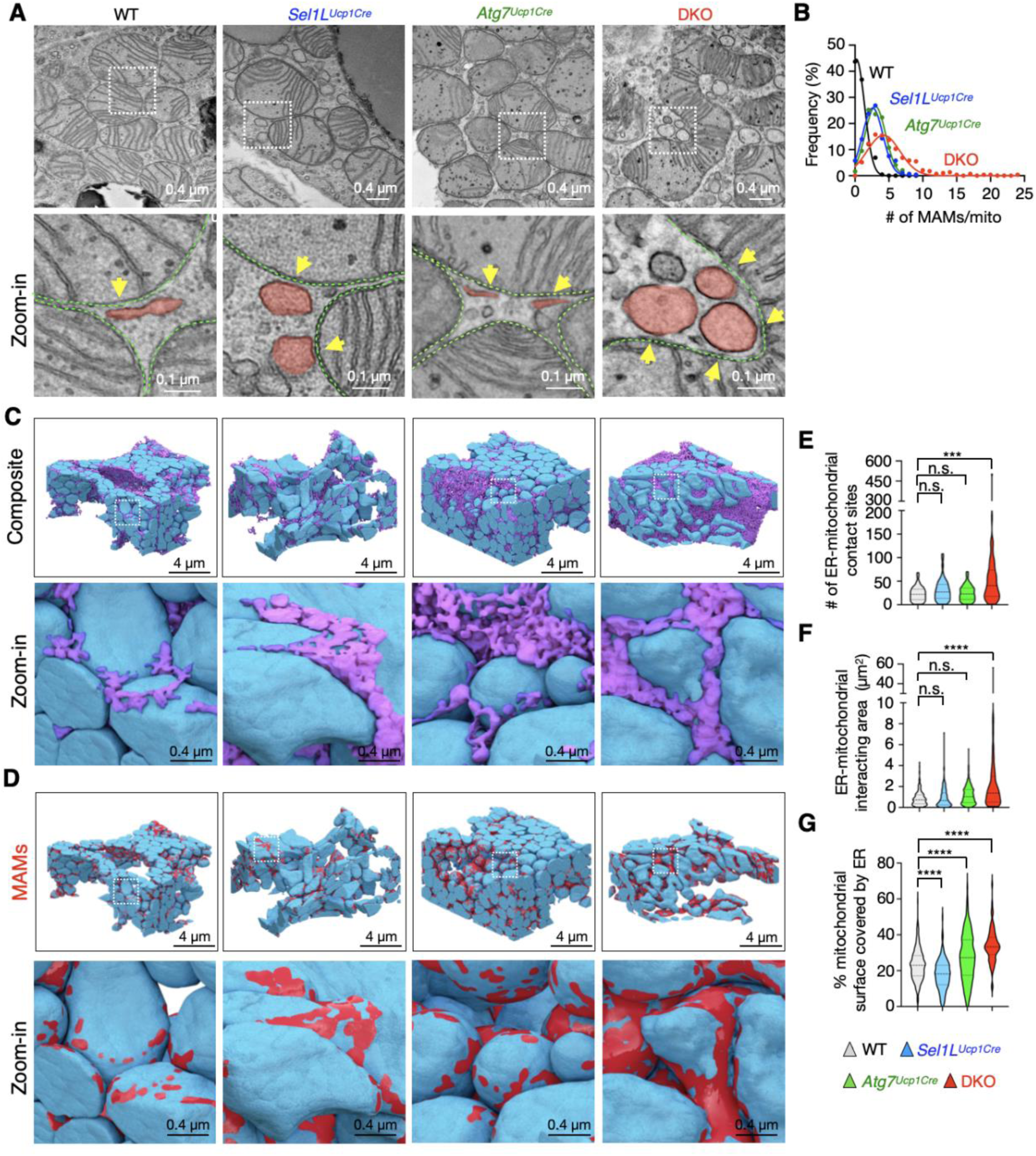
ERAD and autophagy deficiency alters ER morphology and increases ER-mitochondria contacts. (A) Representative TEM images of BAT with ER (red) from 12-week-old male littermates. Arrows point to MAMs; green dotted lines outline mitochondrial membranes. n = 3-5 mice per group. (B) Quantification of number of MAMs per mitochondrion across genotypes. n = 291 mitochondria for WT, 294 for *Sel1L^Ucp1Cre^*, 231 for *Atg7^Ucp1Cre^*, and 242 for DKO, 3-5 mice per group. (C) 3D reconstruction of FIB-SEM images of BAT showing mitochondria (blue) and ER (purple). (D) 3D visualization of mitochondria volumes marked by areas interacting with the ER at 0-25 nm distance (red). (E-G) Quantification of the number of ER-mitochondrial interacting sites per mitochondrion (E), ER-mitochondrial interacting area per mitochondrion (F), and mitochondrion surface covered by ER (G). n = 273 mitochondria for WT, 160 for *Sel1L^Ucp1Cre^*, 375 for *Atg7^Ucp1Cre^*, and 84 for DKO, One-way ANOVA. Data are mean ± SEM. **, p < 0.01; ***, p < 0.001; ****, p < 0.0001.

Three-dimensional FIB-SEM analysis revealed progressive reinforcement of ER–mitochondrial contacts across genotypes, culminating in DKO adipocytes, where mitochondria appeared extensively wrapped by ER membranes (**Figure 5C; Video S1-S4**). To visualize these contacts in 3D, ER voxels were computationally expanded by 25 nm, and interacting surfaces were rendered in red (**Figure 5D; Video S1-S4**). Quantification confirmed a marked increase in the number, area, and extent of ER–mitochondrial interfaces in DKO BAT compared to the other cohorts (**Figure 5E-G**).

To dissect the molecular basis of MAM expansion, we performed subcellular fractionation of BAT to isolate MAMs. The ER–mitochondria tethering proteins vesicle-associated membrane proteins A and B (VAPA/B) were increasingly enriched in the MAM fractions of *Sel1L^Ucp1Cre^* and *Atg7^Ucp1Cre^* BAT, with the highest levels in DKO samples (**Figure S6A**). Notably, SIGMAR1, a known substrate of SEL1L-HRD1 ERAD ^52^, was similarly elevated in MAM fractions of both single KOs and DKO BAT (**Figure S6A**). In contrast, levels of other MAM-associated proteins, including GRP75, VDAC1 (components of the IP3R1-GRP75-VDAC1 complex ^69^), and MFN2, remained largely unchanged (**Figure S6A**). Total protein levels, but not mRNA, of VAPA/B and SIGMAR1 were significantly increased in DKO BAT (**Figure S6B–C**), suggesting post-translational accumulation. These findings suggest that combined loss of SEL1L-HRD1 ERAD and autophagy drives aberrant expansion of MAMs in brown adipocytes, likely through impaired degradation of ER–mitochondria tethering components.

### Mitochondrial hyperfusion and impaired quality control in *DKO* brown adipocytes

To investigate the formation of megamitochondria in DKO brown adipocytes, we examined mitochondrial dynamics. In DKO BAT, both total levels and activating phosphorylation of the mitochondrial fission protein DRP1 were markedly reduced compared to WT controls (**Figure 6A**). In contrast, levels of the outer membrane fusion proteins MFN1 and MFN2 were increased in mitochondria isolated from DKO BAT (**Figure 6B**), despite unchanged total cellular levels across genotypes (**Figure S7A**).

**Figure 6.**
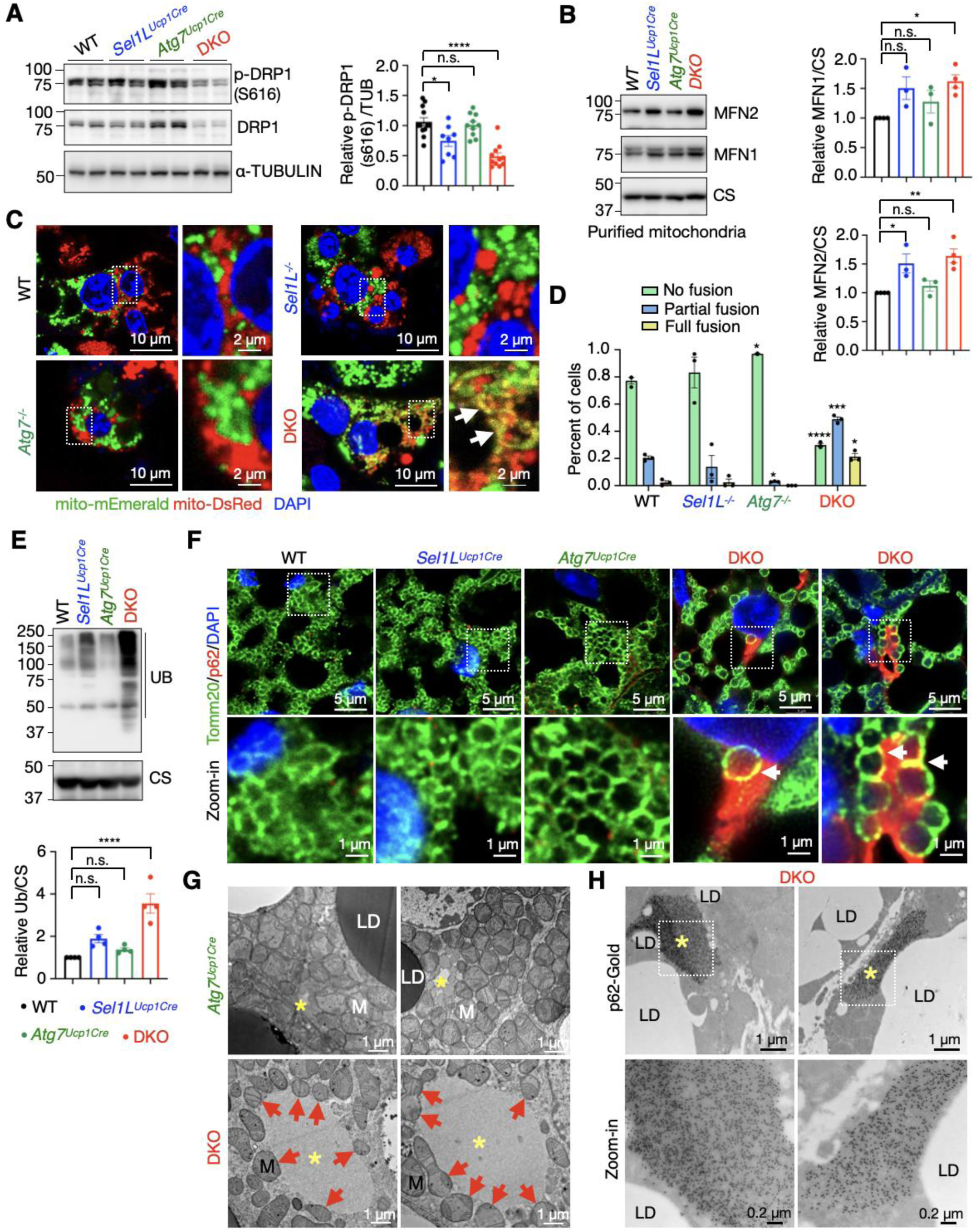
Loss of ERAD and autophagy synergistically promotes mitochondrial fusion, p62 aggregation and mitochondrial quality control. (A) Immunoblot analysis of phosphorylated DRP1 at Ser616 (p-DRP1 s616) as a mitochondrial-fission marker in BAT from 12-week-old littermates, with quantification of p-S616 DRP1 normalized to α-TUBULIN (TUB) shown on the right. n = 8-12 per genotype, One-way ANOVA. (B) Immunoblot analysis of mitochondrial fusion markers mitofusin 1 (MFN1) and MFN2 in isolated mitochondria from BAT of 12-week-old littermates, with quantification of MFN1 and MFN2 levels normalized to citrate synthase (CS) shown on the right. n = 3-4 per genotype, One-way ANOVA. (C-D) Representative confocal images of mitochondrial fusion from the PEG-based fusion assay in mature adipocytes. Mitochondrial fusion (yellow signal) is evidenced by co-localization of mito-DsRed (red) and mito-mEmerald (green). Quantitation of mitochondrial fusion assay shown in (D). Classification of cell hybrids as full, partial, or no fusion. Three independent experiments, with more than 50 cell hybrids from each experiment. Statistical analysis using two-way ANOVA, with comparisons made to the WT group. (E) Immunoblot analysis of ubiquitin (UB) levels in purified mitochondria from BAT. The quantification of UB normalized to citrate synthase (CS) shown below. n = 4, One-way ANOVA. (F) Representative confocal images of TOMM20 (green) and p62 (red) in BAT sections from 12-week-old male littermates. White arrows mark the colocalization of p62 and TOMM20. n = 3 mice per genotype. (G) Representative TEM images of p62 inclusions in BAT from 12-week-old *Atg7^Ucp1Cre^* and DKO male mice. Asterisks indicate p62 inclusion. Red arrows point to mitochondrion near p62 inclusion. LD, lipid droplet; M, mitochondrion. n = 3 mice for *Atg7^Ucp1Cre^* and 5 for DKO. (H) Representative TEM images of BAT following p62-specific immunogold labelling from 12-week-old DKO male mice. Asterisks indicate p62-positive inclusion. n = 2 mice per genotype. Data are mean ± SEM. n.s., not significant; \**p* < 0.1; \*\**p* < 0.01; \*\*\**p* < 0.001; ****, *p* < 0.0001.

To assess fusion activity functionally, we performed a polyethylene glycol (PEG)-based cell fusion assay ^70^ using mature brown adipocytes expressing mitochondrially targeted fluorescent proteins. This assay revealed a significant enhancement in mitochondrial fusion in DKO cells relative to all other genotypes, whereas single KO mice displayed only modest changes (**Figure 6C**, quantified in **Figure 6D**). These findings indicate that simultaneous loss of SEL1L-HRD1 ERAD and autophagy promotes a shift toward excessive mitochondrial fusion.

Notably, hyperfused mitochondria in DKO adipocytes exhibited increased ubiquitination (**Figure 6E**), indicative of elevated mitochondrial damage. Given the established role of p62 in mediating mitophagy ^71,72^, we examined its expression and intracellular distribution. Immunofluorescence staining revealed a marked increase in p62 signal in DKO brown adipocytes, which lacked the typical punctate structures and instead partially aligned along the mitochondrial outer membrane (arrows, **Figure 6F** and **S7B**). Furthermore, TEM showed that cytosolic inclusions – typically formed in the absence of autophagy – were rarely observed in *Atg7^Ucp1Cre^* BAT and completely absent in *Sel1L^Ucp1Cre^* BAT (not shown), but were significantly more numerous and enlarged in DKO adipocytes (asterisks, **Figure 6G** and **S7C**). These inclusions were often closely associated with mitochondria exhibiting disrupted cristae, characteristic of damaged mitochondria (red arrows, **Figure 6G** and **S7C**). Immunogold labeling for p62 further confirmed that these inclusions were p62-positive (asterisks, **Figure 6H**). Together, these findings indicate that the combined loss of ERAD and autophagy synergistically impairs mitophagic clearance, promotes mitochondrial hyperfusion, and leads to the accumulation of ubiquitinated, dysfunctional mitochondria in brown adipocytes.

### Mitochondrial hyperfusion disrupts calcium signaling and impairs thermogenesis in DKO brown adipocytes

To assess the physiological consequences of accumulated megamitochondria, we evaluated calcium dynamics between the ER and mitochondria, a critical process for mitochondrial metabolism and thermogenic function that is mediated by MAMs ^73,74^. Using mito-GCaMP6f, a genetically encoded fluorescent calcium indicator targeted to mitochondria ^75^, we observed that ER-to-mitochondria calcium transfer was significantly reduced in DKO adipocytes compared to *WT* cells (**Figure 7A**). In contrast, ER-to-cytosol calcium release measured with Fluo-4 AM, a cytosolic calcium-sensitive dye ^76^, remained unchanged in DKO adipocytes (**Figure S7A**), suggesting a selective disruption in MAMs-mediated calcium delivery to mitochondria.

**Figure 7.**
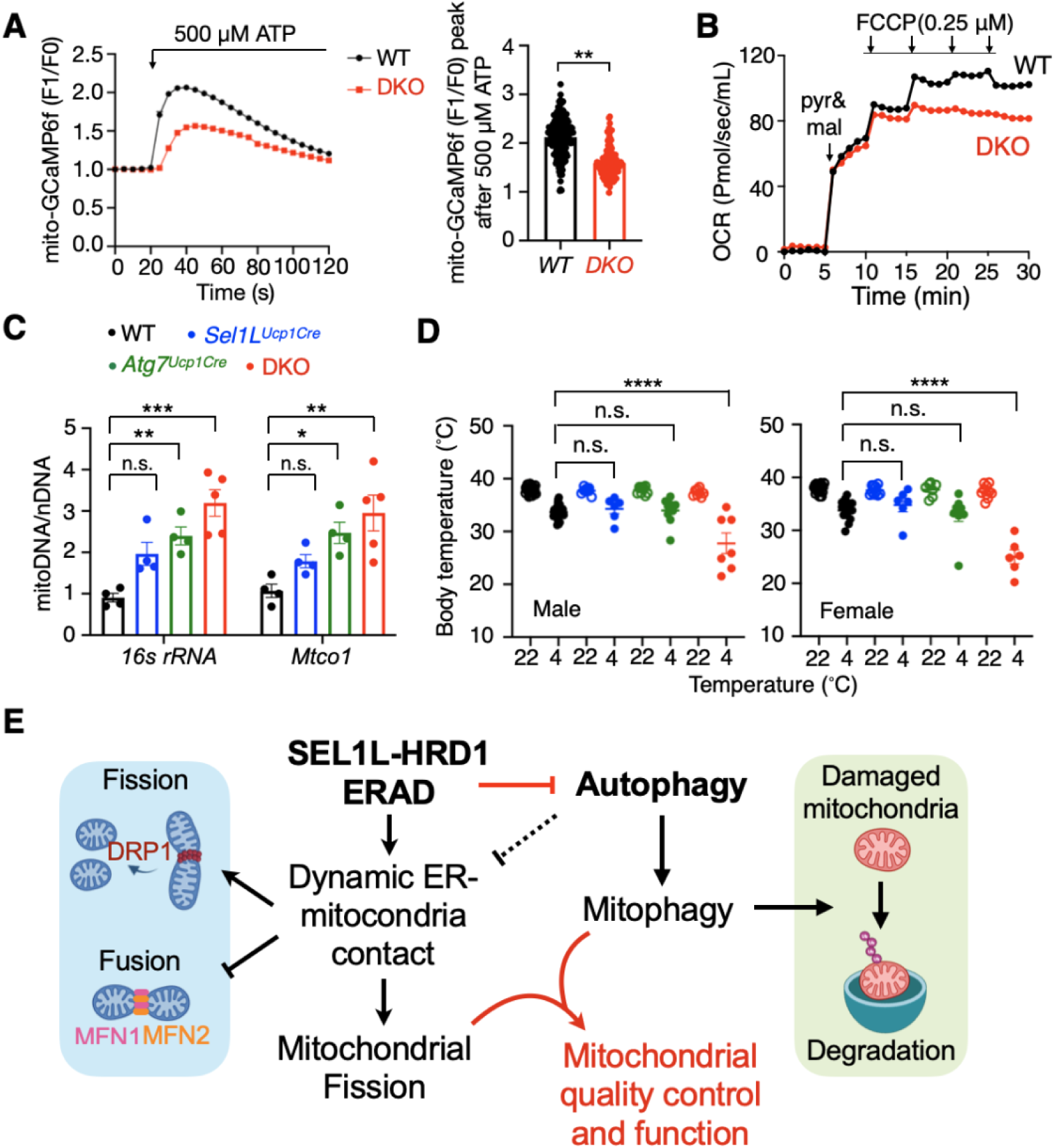
*Sel1L;Atg7* deficiency compromises mitochondria function and thermogenesis in brown adipocytes. (A) Representative traces of mitochondrial Ca^2+^ responds in WT and DKO differentiated adipocytes challenged with 500 μM ATP using mito-GCaMP6f indicator, with quantification of peak mito-GCaMP6f fluorescence shown on the right. n = 177 cells for WT and 129 for DKO, three repeats, Student’s *t*-test. (B) Oxygen consumption rate (OCR) of purified mitochondria from BAT of 12-week-old WT and DKO male littermates. Three repeats. (C) qPCR analyses of mitochondrial DNA targeting mitochondrial 1*6s rRNA* and *Mtco1* genes, normalized to the nuclear-encoded *Hk2* gene. n = 4-5 per group, One-way ANOVA. (D) Rectal temperatures of 12-week-old male and female littermates exposed to 4°C for 2 hours. n = 6-21 per group, Two-way ANOVA. (E) Proposed model for a synergic role of SEL1L-HRD1 ERAD and autophagy in mitochondrial quality control in brown adipocytes. See text for details. Data are mean ± SEM. n.s., not significant; \**p* < 0.1; \*\**p* < 0.01; \*\*\**p* < 0.001; ****, *p* < 0.0001.

To determine the impact on mitochondrial respiration, we measured oxygen consumption rates in mitochondria isolated from BAT. Purified DKO mitochondria exhibited a markedly reduced maximal respiratory capacity in response to FCCP, comparable to the reduction observed in *Sel1L^Ucp1Cre^* mice but contrasting with enhanced respiration seen in *Atg7^Ucp1Cre^* mitochondria (**Figure 7B** and **S8B-C**). Blue-native PAGE analysis of oxidative phosphorylation (OXPHOS) complexes revealed diminished abundance of complexes III and V in both *Sel1L^Ucp1Cre^* and DKO BAT, despite unchanged levels of individual complex subunits by immunoblotting (**Figure S8D** and quantitated in **Fig. S8E**). Consistent with compromised mitochondrial function, DKO BAT displayed elevated mitochondrial DNA (mtDNA) content (**Figure 7C**), suggestive of compensatory mitochondrial biogenesis.

Lastly, we assessed thermogenic capacity under cold stress. At room temperature, body temperatures were comparable across genotypes. However, during acute cold exposure, DKO mice rapidly developed hypothermia, exhibiting a significant drop in core body temperature within 2 hours (**Figure 7D**). In contrast, WT and single-KO mice maintained euthermia over this interval. Notably, *Sel1L^Ucp1Cre^* mice only became hypothermic after prolonged cold exposure (4–6 hours), as previously reported ^52^. Together, these findings highlight the cooperative role of SEL1L-HRD1 ERAD and autophagy in preserving mitochondrial calcium homeostasis, respiratory competence, and thermogenic function in brown adipocytes.

## DISCUSSION

Our study uncovers a previously unrecognized interplay between SEL1L-HRD1 ER-associated degradation (ERAD) and autophagy in the maintenance of mitochondrial quality control and function in brown adipocytes. In the absence of *Sel1L*, compensatory autophagy is activated, supporting both ER homeostasis and mitochondrial integrity. However, the combined deficiency of ERAD and autophagy leads to the formation of pleomorphic “megamitochondria,” impaired mitochondrial respiration, and severe thermogenic dysfunction (Figure 7E). These megamitochondria are characterized by increased ER-mitochondria contacts and elevated levels of MAM proteins, such as VAPA/B and SIGMAR1. Given the established role of these contacts in mitochondrial fission ^54^, our findings suggest that these alterations disrupt mitochondrial dynamics by inhibiting fission and promoting fusion. This is evident even at room temperature, where megamitochondria are perforated by ER tubules. Notably, these structural and functional abnormalities are absent in brown adipocytes of single KO ^52^ or *Sel1L*-*Ire1a* double-deficient mice housed at room temperature, highlighting the distinct and synergistic roles of ERAD and autophagy in regulating mitochondrial dynamics. Collectively, our findings demonstrate the integrated roles of ERAD and autophagy in preserving ER and mitochondrial integrity, their interactions, and mitochondrial function.

The ERAD-autophagy interaction in brown adipocytes is distinct from its role in white adipocytes and pancreatic β cells, where these pathways primarily collaborate to clear protein aggregates, such as lipoprotein lipase and proinsulin, ensuring ER homeostasis without noticeable impact on mitochondrial morphology ^62,63^. In brown adipocytes, however, the effect extends to mitochondria, likely due to the high mitochondrial content and dynamic nature of mitochondria. Previously, we demonstrated that ERAD regulates ER-mitochondrial contacts via MAM proteins, with megamitochondria only observed in *Sel1L*-deficient mice after 4-6 hours of cold exposure ^52^. In contrast, here we show that megamitochondria formation in ERAD-autophagy DKO cells occurs even at room temperature, independent of cold exposure. We speculate that this synergistic effect depends not only on ER expansion and the accumulation of damaged mitochondria, but also from increased ER-mitochondria contacts and elevated MAM protein levels, which likely stall mitochondrial fission.

Our finding of increased VAPA/B protein abundance at MAMs in cells deficient in both ERAD and autophagy adds a new dimension to the understanding of ER-mitochondrial dynamics. VAPA/B is a key ER receptor involved in interactions with other organelles, such as the Golgi, mitochondria, and endosomes/autophagosomes ^77–79^, facilitating lipid exchange and maintaining membrane integrity. The elevated levels of VAPA/B may reflect a cellular attempt to enhance lipid transfer between organelles under stress, given their critical role in organelle function. However, this adaptive response likely has both beneficial and detrimental effects. While increased ER-mitochondrial contacts may temporarily support mitochondrial lipid requirements, the resulting mitochondrial hyperfusion—exacerbated by the accumulation of damaged mitochondria due to autophagy deficiency—may impair respiratory function ^80^. This highlights the critical balance between mitochondrial fusion and fission, which is critical for mitochondrial health. Dysregulation of this balance, in the context of combined ERAD and autophagy deficiencies, likely contributes significantly to mitochondrial dysfunction.

The impaired mitochondrial respiration and thermogenic responses observed in our study highlight the broader metabolic consequences of combined SEL1L-HRD1 ERAD and autophagy deficiencies. As brown adipose tissue (BAT) plays a central role in energy expenditure and thermogenesis in humans ^81,82^, and considering that mitochondrial function and dynamics are critical to the pathogenesis of various diseases ^83,84^, our findings have significant implications for metabolic disorders such as obesity and type 2 diabetes, as well as for neurodegeneration and aging. These results suggest that the interplay between ERAD and autophagy in mitochondrial quality control could serve as a promising therapeutic target, especially in light of our recent identification of patients with hypomorphic ERAD mutations ^50,51^. Enhancing autophagic activity in ERAD-deficient cells may restore mitochondrial function and improve metabolic outcomes. Further investigation into the molecular mechanisms driving ERAD-autophagy synergy is crucial for developing therapeutic strategies to maintain mitochondrial integrity and address metabolic dysfunction. Future studies should focus on translating these findings into actionable treatments to improve mitochondrial function and metabolic health.

### Limitations of the Study

Our study identifies a synergistic role for SEL1L-HRD1 ERAD and autophagy in maintaining mitochondrial quality control in brown adipocytes. However, two limitations should be acknowledged. First, although we observed enhanced ER-mitochondria contacts and increased expression of MAM-associated tethering proteins in DKO BAT, we did not directly establish a causal relationship between altered MAM dynamics and mitochondrial hyperfusion. Further mechanistic studies, such as manipulating specific MAM components, will be necessary to delineate their direct contribution to mitochondrial remodeling. Second, our use of the Ucp1-Cre driver to generate adipocyte-specific knockouts may introduce unintended effects, as Ucp1-Cre is known to exhibit off-target recombination in certain non-adipose tissues, including the central nervous system and skeletal muscle. This could confound the interpretation of brown adipocyte-specific phenotypes, especially for systemic or whole-tissue analyses.

## Resource availability

### Lead contact

Further information and requests for resources and reagents should be directed to and will be fulfilled by the lead contact, Ling Qi.

### Materials Availability

All unique/stable reagents generated in this study are available from the lead contact without restriction.

### Data and Code Availability

This study did not generate large-scale datasets or original code. All data supporting the findings of this study are included in the main text or Supplemental Materials and are available from the Lead Contact upon reasonable request.

## ACKNOWLEDGEMENTS

We acknowledge Drs. Ming-Feng Tsai and Yanzhuang Wang for providing reagents; Allen Hunter for assistance with FIB-SEM analysis; Dave Castle, members of the Qi–Sun and Arvan laboratories for technical assistance and insightful discussions. We acknowledge the use of Electron Microscopy Core, Advanced Microscopy Core, Research Histology Core at the University of Virginia, and the University of Michigan Comprehensive Cancer Center (UMCCC) Tissue Core, Imaging laboratory, the Microscopy Core of Michigan Biomedical Research Core Facilities, and Michigan Center for Materials Characterization. This work was supported by NIH grants 1R01DK11174, R01DK117639, R35GM130292 and 1R01DK120047 (L.Q.). X.C. is supported by American Diabetes Association (ADA) Postdoctoral Fellowship (11-23-PDF-62).

## AUTHOR CONTRIBUTIONS

X.C. and S.W. collaboratively designed and performed most experiments; M.T. assisted with EM experiments. L.Q. and S.S. supervised the study and wrote the manuscript with help from X.C. X.C. wrote the methods and figure legends; S.H. provided technical support for data analysis. All authors commented on and approved the final manuscript.

## REFERENCES

1. van der Bliek, A.M., Shen, Q., and Kawajiri, S. (2013). Mechanisms of mitochondrial fission and fusion. Cold Spring Harb Perspect Biol 5. 10.1101/cshperspect.a011072.

2. Pickles, S., Vigie, P., and Youle, R.J. (2018). Mitophagy and Quality Control Mechanisms in Mitochondrial Maintenance. Curr Biol 28, R170–R185. 10.1016/j.cub.2018.01.004.

3. Aquilano, K., Zhou, B., Brestoff, J.R., and Lettieri-Barbato, D. (2023). Multifaceted mitochondrial quality control in brown adipose tissue. Trends Cell Biol 33, 517–529. 10.1016/j.tcb.2022.09.008.

4. Oelkrug, R., Polymeropoulos, E.T., and Jastroch, M. (2015). Brown adipose tissue: physiological function and evolutionary significance. J Comp Physiol B 185, 587–606. 10.1007/s00360-015-0907-7.

5. Wikstrom, J.D., Mahdaviani, K., Liesa, M., Sereda, S.B., Si, Y., Las, G., Twig, G., Petrovic, N., Zingaretti, C., Graham, A., et al. (2014). Hormone-induced mitochondrial fission is utilized by brown adipocytes as an amplification pathway for energy expenditure. EMBO J 33, 418–436. 10.1002/embj.201385014.

6. Lu, Y., Fujioka, H., Joshi, D., Li, Q., Sangwung, P., Hsieh, P., Zhu, J., Torio, J., Sweet, D., Wang, L., et al. (2018). Mitophagy is required for brown adipose tissue mitochondrial homeostasis during cold challenge. Sci Rep 8, 8251. 10.1038/s41598-018-26394-5.

7. Rosina, M., Ceci, V., Turchi, R., Chuan, L., Borcherding, N., Sciarretta, F., Sanchez-Diaz, M., Tortolici, F., Karlinsey, K., Chiurchiu, V., et al. (2022). Ejection of damaged mitochondria and their removal by macrophages ensure efficient thermogenesis in brown adipose tissue. Cell Metab 34, 533–548 e512. 10.1016/j.cmet.2022.02.016.

8. Mizushima, N., Noda, T., Yoshimori, T., Tanaka, Y., Ishii, T., George, M.D., Klionsky, D.J., Ohsumi, M., and Ohsumi, Y. (1998). A protein conjugation system essential for autophagy. Nature 395, 395–398. 10.1038/26506.

9. Choi, A.M., Ryter, S.W., and Levine, B. (2013). Autophagy in human health and disease. N Engl J Med 368, 651–662. 10.1056/NEJMra1205406.

10. Levine, B., and Kroemer, G. (2008). Autophagy in the pathogenesis of disease. Cell 132, 27–42. 10.1016/j.cell.2007.12.018.

11. Martinez-Lopez, N., and Singh, R. (2015). Autophagy and Lipid Droplets in the Liver. Annu Rev Nutr 35, 215–237. 10.1146/annurev-nutr-071813-105336.

12. Hwang, J., and Qi, L. (2018). Quality Control in the Endoplasmic Reticulum: Crosstalk between ERAD and UPR pathways. Trends Biochem Sci 43, 593–605. 10.1016/j.tibs.2018.06.005.

13. Bhattacharya, A., and Qi, L. (2019). ER-associated degradation in health and disease - from substrate to organism. J Cell Sci 132, jcs232850. 10.1242/jcs.232850.

14. Wu, S.A., Li, Z.J., and Qi, L. (2025). Endoplasmic reticulum (ER) protein degradation by ER-associated degradation and ER-phagy. Trends Cell Biol, S0962-8924(0925)00002-00009. 10.1016/j.tcb.2025.01.002.

15. Yang, Z., and Klionsky, D.J. (2010). Eaten alive: a history of macroautophagy. Nat Cell Biol 12, 814–822. 10.1038/ncb0910-814.

16. Kroemer, G., Marino, G., and Levine, B. (2010). Autophagy and the integrated stress response. Mol Cell 40, 280–293. 10.1016/j.molcel.2010.09.023.

17. Hubner, C.A., and Dikic, I. (2020). ER-phagy and human diseases. Cell Death Differ 27, 833–842. 10.1038/s41418-019-0444-0.

18. Molinari, M. (2021). ER-phagy responses in yeast, plants, and mammalian cells and their crosstalk with UPR and ERAD. Dev Cell 56, 949–966. 10.1016/j.devcel.2021.03.005.

19. Bernales, S., Schuck, S., and Walter, P. (2007). ER-phagy: selective autophagy of the endoplasmic reticulum. Autophagy 3, 285–287. 10.4161/auto.3930.

20. Onishi, M., Yamano, K., Sato, M., Matsuda, N., and Okamoto, K. (2021). Molecular mechanisms and physiological functions of mitophagy. EMBO J 40, e104705. 10.15252/embj.2020104705.

21. Levine, B., and Kroemer, G. (2019). Biological Functions of Autophagy Genes: A Disease Perspective. Cell 176, 11–42. 10.1016/j.cell.2018.09.048.

22. Singh, R., Xiang, Y., Wang, Y., Baikati, K., Cuervo, A.M., Luu, Y.K., Tang, Y., Pessin, J.E., Schwartz, G.J., and Czaja, M.J. (2009). Autophagy regulates adipose mass and differentiation in mice. J Clin Invest 119, 3329–3339. 10.1172/JCI39228.

23. Zhang, Y., Goldman, S., Baerga, R., Zhao, Y., Komatsu, M., and Jin, S. (2009). Adipose-specific deletion of autophagy-related gene 7 (atg7) in mice reveals a role in adipogenesis. Proc Natl Acad Sci U S A 106, 19860–19865. 10.1073/pnas.0906048106.

24. Altshuler-Keylin, S., Shinoda, K., Hasegawa, Y., Ikeda, K., Hong, H., Kang, Q., Yang, Y., Perera, R.M., Debnath, J., and Kajimura, S. (2016). Beige Adipocyte Maintenance Is Regulated by Autophagy-Induced Mitochondrial Clearance. Cell Metab 24, 402–419. 10.1016/j.cmet.2016.08.002.

25. Cai, J., Pires, K.M., Ferhat, M., Chaurasia, B., Buffolo, M.A., Smalling, R., Sargsyan, A., Atkinson, D.L., Summers, S.A., Graham, T.E., and Boudina, S. (2018). Autophagy Ablation in Adipocytes Induces Insulin Resistance and Reveals Roles for Lipid Peroxide and Nrf2 Signaling in Adipose-Liver Crosstalk. Cell reports 25, 1708–1717 e1705. 10.1016/j.celrep.2018.10.040.

26. Lippincott-Schwartz, J., Bonifacino, J.S., Yuan, L.C., and Klausner, R.D. (1988). Degradation from the endoplasmic reticulum: disposing of newly synthesized proteins. Cell 54, 209–220. 10.1016/0092-8674(88)90553-3.

27. Sommer, T., and Jentsch, S. (1993). A protein translocation defect linked to ubiquitin conjugation at the endoplasmic reticulum. Nature 365, 176–179. 10.1038/365176a0.

28. Jensen, T.J., Loo, M.A., Pind, S., Williams, D.B., Goldberg, A.L., and Riordan, J.R. (1995). Multiple proteolytic systems, including the proteasome, contribute to CFTR processing. Cell 83, 129–135. 10.1016/0092-8674(95)90241-4.

29. Ward, C.L., Omura, S., and Kopito, R.R. (1995). Degradation of CFTR by the ubiquitin-proteasome pathway. Cell 83, 121–127.

30. McCracken, A.A., and Brodsky, J.L. (1996). Assembly of ER-associated protein degradation in vitro: dependence on cytosol, calnexin, and ATP. J Cell Biol 132, 291–298. 10.1083/jcb.132.3.291.

31. Hampton, R.Y., Gardner, R.G., and Rine, J. (1996). Role of 26S proteasome and HRD genes in the degradation of 3-hydroxy-3-methylglutaryl-CoA reductase, an integral endoplasmic reticulum membrane protein. Mol Biol Cell 7, 2029–2044. 10.1091/mbc.7.12.2029.

32. Guerriero, C.J., and Brodsky, J.L. (2012). The delicate balance between secreted protein folding and endoplasmic reticulum-associated degradation in human physiology. Physiol Rev 92, 537–576. 10.1152/physrev.00027.2011.

33. Sun, Z., and Brodsky, J.L. (2019). Protein quality control in the secretory pathway. J Cell Biol 218, 3171–3187. 10.1083/jcb.201906047.

34. Christianson, J.C., Jarosch, E., and Sommer, T. (2023). Mechanisms of substrate processing during ER-associated protein degradation. Nat Rev Mol Cell Biol 24, 777–796. 10.1038/s41580-023-00633-8.

35. Mueller, B., Lilley, B.N., and Ploegh, H.L. (2006). SEL1L, the homologue of yeast Hrd3p, is involved in protein dislocation from the mammalian ER. J Cell Biol 175, 261–270. jcb.200605196 [pii]10.1083/jcb.200605196.

36. Mueller, B., Klemm, E.J., Spooner, E., Claessen, J.H., and Ploegh, H.L. (2008). SEL1L nucleates a protein complex required for dislocation of misfolded glycoproteins. Proc Natl Acad Sci USA 105, 12325–12330. 10.1073/pnas.0805371105.

37. Plemper, R.K., Bordallo, J., Deak, P.M., Taxis, C., Hitt, R., and Wolf, D.H. (1999). Genetic interactions of Hrd3p and Der3p/Hrd1p with Sec61p suggest a retro-translocation complex mediating protein transport for ER degradation. J Cell Sci 112 ( *Pt 22*), 4123–4134.

38. Sun, S., Shi, G., Han, X., Francisco, A.B., Ji, Y., Mendonca, N., Liu, X., Locasale, J.W., Simpson, K.W., Duhamel, G.E., et al. (2014). Sel1L is indispensable for mammalian endoplasmic reticulum-associated degradation, endoplasmic reticulum homeostasis, and survival. Proc Natl Acad Sci U S A 111, E582–591. 10.1073/pnas.1318114111.

39. Lin, L.L., Wang, H.H., Pederson, B., Wei, X., Torres, M., Lu, Y., Li, Z.J., Liu, X., Mao, H., Wang, H., et al. (2024). SEL1L-HRD1 interaction is required to form a functional HRD1 ERAD complex. Nature communications 15, 1440. 10.1038/s41467-024-45633-0.

40. Francisco, A.B., Singh, R., Li, S., Vani, A.K., Yang, L., Munroe, R.J., Diaferia, G., Cardano, M., Biunno, I., Qi, L., et al. (2010). Deficiency of suppressor enhancer lin12 1 like (SEL1L) in mice leads to systemic endoplasmic reticulum stress and embryonic lethality. J Biol Chem 285, 13694–13703. M109.085340 [pii] 10.1074/jbc.M109.085340.

41. Yagishita, N., Ohneda, K., Amano, T., Yamasaki, S., Sugiura, A., Tsuchimochi, K., Shin, H., Kawahara, K., Ohneda, O., Ohta, T., et al. (2005). Essential role of synoviolin in embryogenesis. J Biol Chem 280, 7909–7916. 10.1074/jbc.M410863200.

42. Fujita, H., Yagishita, N., Aratani, S., Saito-Fujita, T., Morota, S., Yamano, Y., Hansson, M.J., Inazu, M., Kokuba, H., Sudo, K., et al. (2015). The E3 ligase synoviolin controls body weight and mitochondrial biogenesis through negative regulation of PGC-1beta. EMBO J 34, 1042–1055. 10.15252/embj.201489897.

43. Qi, L., Tsai, B., and Arvan, P. (2017). New Insights into the Physiological Role of Endoplasmic Reticulum-Associated Degradation. Trends Cell Biol 27, 430–440. 10.1016/j.tcb.2016.12.002.

44. Sha, H., Sun, S., Francisco, A.B., Ehrhardt, N., Xue, Z., Liu, L., Lawrence, P., Mattijssen, F., Guber, R.D., Panhwar, M.S., et al. (2014). The ER-associated degradation adaptor protein Sel1L regulates LPL secretion and lipid metabolism. Cell Metab 20, 458–470.

45. Ji, Y., Kim, H., Yang, L., Sha, H., Roman, C.A., Long, Q., and Qi, L. (2016). The Sel1L-Hrd1 Endoplasmic Reticulum-Associated Degradation Complex Manages a Key Checkpoint in B Cell Development. Cell reports 16, 2630–2640. 10.1016/j.celrep.2016.08.003.

46. Sun, S., Louri, R., Cohen, S.B., Ji, Y., Goodrich, J.K., Poole, A.C., Ley, R.E., Denkers, E.Y., Mcguckin, M.A., Long, Q., et al. (2016). Epithelial Sel1L is required for the maintenance of intestinal homeostasis. Mol Biol Cell 27, 483–490.

47. Yoshida, S., Wei, X., Zhang, G., O’Connor, C.L., Torres, M., Zhou, Z., Lin, L., Menon, R., Xu, X., Zheng, W., et al. (2021). Endoplasmic reticulum-associated degradation is required for nephrin maturation and kidney glomerular filtration function. J Clin Invest 131. 10.1172/JCI143988.

48. Shi, G., Somlo, D., Kim, G.H., Prescianotto-Baschong, C., Sun, S., Beuret, N., Long, Q., Rutishauser, J., Arvan, P., Spiess, M., and Qi, L. (2017). ER-associated degradation is required for vasopressin prohormone processing and systemic water homeostasis. J Clin Invest 127, 3897–3912.

49. Kim, G.H., Shi, G., Somlo, D.R.M., Haataja, L., Soon, S., Long, Q., Nillni, E.A., Low, M.J., Arvan, P., Myers, M.G., and Qi, L. (2018). Hypothalamic ER-associated degradation regulates POMC maturation, feeding and age-associated obesity. J Clin Invest 128, 1125–1140.

50. Wang, H.H., Lin, L.L., Li, Z.J., Wei, X., Askander, O., Cappuccio, G., Hashem, M.O., Hubert, L., Munnich, A., Alqahtani, M., et al. (2024). Hypomorphic variants of SEL1L-HRD1 ER-associated degradation are associated with neurodevelopmental disorders. J Clin Invest 134. 10.1172/JCI170054.

51. Weis, D., Lin, L.L., Wang, H.H., Li, Z.J., Kusikova, K., Ciznar, P., Wolf, H.M., Leiss-Piller, A., Wang, Z., Wei, X., et al. (2024). Biallelic Cys141Tyr variant of SEL1L is associated with neurodevelopmental disorders, agammaglobulinemia, and premature death. J Clin Invest 134. 10.1172/JCI170882.

52. Zhou, Z., Torres, M., Sha, H., Halbrook, C.J., Van den Bergh, F., Reinert, R.B., Yamada, T., Wang, S., Luo, Y., Hunter, A.H., et al. (2020). Endoplasmic reticulum-associated degradation regulates mitochondrial dynamics in brown adipocytes. Science 368, 54–60. 10.1126/science.aay2494.

53. Koch, C., Schuldiner, M., and Herrmann, J.M. (2021). ER-SURF: Riding the Endoplasmic Reticulum Surface to Mitochondria. Int J Mol Sci 22. 10.3390/ijms22179655.

54. Friedman, J.R., Lackner, L.L., West, M., DiBenedetto, J.R., Nunnari, J., and Voeltz, G.K. (2011). ER tubules mark sites of mitochondrial division. Science 334, 358–362. 10.1126/science.1207385.

55. Rowland, A.A., and Voeltz, G.K. (2012). Endoplasmic reticulum-mitochondria contacts: function of the junction. Nat Rev Mol Cell Biol 13, 607–625. 10.1038/nrm3440.

56. Wu, H., Carvalho, P., and Voeltz, G.K. (2018). Here, there, and everywhere: The importance of ER membrane contact sites. Science 361. 10.1126/science.aan5835.

57. Rusinol, A.E., Cui, Z., Chen, M.H., and Vance, J.E. (1994). A unique mitochondria-associated membrane fraction from rat liver has a high capacity for lipid synthesis and contains pre-Golgi secretory proteins including nascent lipoproteins. J Biol Chem 269, 27494–27502.

58. Vance, J.E. (2014). MAM (mitochondria-associated membranes) in mammalian cells: lipids and beyond. Biochim Biophys Acta 1841, 595–609. 10.1016/j.bbalip.2013.11.014.

59. Murley, A., Lackner, L.L., Osman, C., West, M., Voeltz, G.K., Walter, P., and Nunnari, J. (2013). ER-associated mitochondrial division links the distribution of mitochondria and mitochondrial DNA in yeast. eLife 2, e00422. 10.7554/eLife.00422.

60. Lewis, S.C., Uchiyama, L.F., and Nunnari, J. (2016). ER-mitochondria contacts couple mtDNA synthesis with mitochondrial division in human cells. Science 353, aaf5549. 10.1126/science.aaf5549.

61. Youle, R.J., and Narendra, D.P. (2011). Mechanisms of mitophagy. Nat Rev Mol Cell Biol 12, 9–14. 10.1038/nrm3028.

62. Shrestha, N., Torres, M., Zhang, J., Lu, Y., Haataja, L., Reinert, R.B., Knupp, J., Chen, Y.J., Parlakgul, G., Arruda, A.P., et al. (2023). Integration of ER protein quality control mechanisms defines beta cell function and ER architecture. J Clin Invest 133, e163584. 10.1172/JCI163584.

63. Wu, S.A., Shen, C., Wei, X., Zhang, X., Wang, S., Chen, X., Torres, M., Lu, Y., Lin, L.L., Wang, H.H., et al. (2023). The mechanisms to dispose of misfolded proteins in the endoplasmic reticulum of adipocytes. Nature communications 14, 3132. 10.1038/s41467-023-38690-4.

64. Walter, P., and Ron, D. (2011). The unfolded protein response: from stress pathway to homeostatic regulation. Science 334, 1081–1086. 10.1126/science.1209038.

65. Claflin, K.E., Flippo, K.H., Sullivan, A.I., Naber, M.C., Zhou, B., Neff, T.J., Jensen-Cody, S.O., and Potthoff, M.J. (2022). Conditional gene targeting using UCP1-Cre mice directly targets the central nervous system beyond thermogenic adipose tissues. Molecular metabolism 55, 101405. 10.1016/j.molmet.2021.101405.

66. Sun, S., Shi, G., Sha, H., Ji, Y., Han, X., Shu, X., Ma, H., Inoue, T., Gao, B., Kim, H., et al. (2015). IRE1a is an endogenous substrate of endoplasmic-reticulum-associated degradation. Nat Cell Biol 17, 1546–1555. 10.1038/ncb3266.

67. Flis, V.V., and Daum, G. (2013). Lipid transport between the endoplasmic reticulum and mitochondria. Cold Spring Harbor perspectives in biology 5. 10.1101/cshperspect.a013235.

68. Khaminets, A., Heinrich, T., Mari, M., Grumati, P., Huebner, A.K., Akutsu, M., Liebmann, L., Stolz, A., Nietzsche, S., Koch, N., et al. (2015). Regulation of endoplasmic reticulum turnover by selective autophagy. Nature 522, 354–358. 10.1038/nature14498.

69. Szabadkai, G., Bianchi, K., Varnai, P., De Stefani, D., Wieckowski, M.R., Cavagna, D., Nagy, A.I., Balla, T., and Rizzuto, R. (2006). Chaperone-mediated coupling of endoplasmic reticulum and mitochondrial Ca2+ channels. J Cell Biol 175, 901–911. 10.1083/jcb.200608073.

70. Nascimento, A., Lannigan, J., and Kashatus, D. (2016). High-throughput detection and quantification of mitochondrial fusion through imaging flow cytometry. Cytometry A 89, 708–719. 10.1002/cyto.a.22891.

71. Yamada, T., Murata, D., Adachi, Y., Itoh, K., Kameoka, S., Igarashi, A., Kato, T., Araki, Y., Huganir, R.L., Dawson, T.M., et al. (2018). Mitochondrial Stasis Reveals p62-Mediated Ubiquitination in Parkin-Independent Mitophagy and Mitigates Nonalcoholic Fatty Liver Disease. Cell Metab 28, 588–604 e585. 10.1016/j.cmet.2018.06.014.

72. Yamada, T., Dawson, T.M., Yanagawa, T., Iijima, M., and Sesaki, H. (2019). SQSTM1/p62 promotes mitochondrial ubiquitination independently of PINK1 and PRKN/parkin in mitophagy. Autophagy 15, 2012–2018. 10.1080/15548627.2019.1643185.

73. Giorgi, C., Marchi, S., and Pinton, P. (2018). Publisher Correction: The machineries, regulation and cellular functions of mitochondrial calcium. Nat Rev Mol Cell Biol 19, 746. 10.1038/s41580-018-0066-2.

74. Sassano, M.L., Felipe-Abrio, B., and Agostinis, P. (2022). ER-mitochondria contact sites; a multifaceted factory for Ca(2+) signaling and lipid transport. Front Cell Dev Biol 10, 988014. 10.3389/fcell.2022.988014.

75. Chen, T.W., Wardill, T.J., Sun, Y., Pulver, S.R., Renninger, S.L., Baohan, A., Schreiter, E.R., Kerr, R.A., Orger, M.B., Jayaraman, V., et al. (2013). Ultrasensitive fluorescent proteins for imaging neuronal activity. Nature 499, 295–300. 10.1038/nature12354.

76. Gee, K.R., Brown, K.A., Chen, W.N., Bishop-Stewart, J., Gray, D., and Johnson, I. (2000). Chemical and physiological characterization of fluo-4 Ca(2+)-indicator dyes. Cell Calcium 27, 97–106. 10.1054/ceca.1999.0095.

77. Dong, R., Saheki, Y., Swarup, S., Lucast, L., Harper, J.W., and De Camilli, P. (2016). Endosome-ER Contacts Control Actin Nucleation and Retromer Function through VAP-Dependent Regulation of PI4P. Cell 166, 408–423. 10.1016/j.cell.2016.06.037.

78. Zhao, Y.G., Liu, N., Miao, G., Chen, Y., Zhao, H., and Zhang, H. (2018). The ER Contact Proteins VAPA/B Interact with Multiple Autophagy Proteins to Modulate Autophagosome Biogenesis. Curr Biol 28, 1234–1245 e1234. 10.1016/j.cub.2018.03.002.

79. Mao, D., Lin, G., Tepe, B., Zuo, Z., Tan, K.L., Senturk, M., Zhang, S., Arenkiel, B.R., Sardiello, M., and Bellen, H.J. (2019). VAMP associated proteins are required for autophagic and lysosomal degradation by promoting a PtdIns4P-mediated endosomal pathway. Autophagy 15, 1214–1233. 10.1080/15548627.2019.1580103.

80. Wai, T., and Langer, T. (2016). Mitochondrial Dynamics and Metabolic Regulation. Trends Endocrinol Metab 27, 105–117. 10.1016/j.tem.2015.12.001.

81. Leitner, B.P., Huang, S., Brychta, R.J., Duckworth, C.J., Baskin, A.S., McGehee, S., Tal, I., Dieckmann, W., Gupta, G., Kolodny, G.M., et al. (2017). Mapping of human brown adipose tissue in lean and obese young men. Proc Natl Acad Sci U S A 114, 8649–8654. 10.1073/pnas.1705287114.

82. Porter, C., Herndon, D.N., Chondronikola, M., Chao, T., Annamalai, P., Bhattarai, N., Saraf, M.K., Capek, K.D., Reidy, P.T., Daquinag, A.C., et al. (2016). Human and Mouse Brown Adipose Tissue Mitochondria Have Comparable UCP1 Function. Cell Metab 24, 246–255. 10.1016/j.cmet.2016.07.004.

83. Archer, S.L. (2013). Mitochondrial dynamics--mitochondrial fission and fusion in human diseases. N Engl J Med 369, 2236–2251. 10.1056/NEJMra1215233.

84. Chen, H., and Chan, D.C. (2009). Mitochondrial dynamics--fusion, fission, movement, and mitophagy--in neurodegenerative diseases. Hum Mol Genet 18, R169–176. 10.1093/hmg/ddp326.

