## Supplemental information for "Synergistic Control of Mitochondrial Dynamics and Function by ERAD and Autophagy in Brown Adipocytes"

### STAR★METHODS

#### Key resources table

| REAGENT or RESOURCE | SOURCE | IDENTIFIER |
|---|---|---|
| Antibodies | | |
| Rabbit monoclonal anti-Cleaved Caspase-3 (Asp175) | Cell Signaling Technology | Cat# 9661; RRID: AB_2341188 |
| Mouse monoclonal anti-HSP90α/β (H-114) | Santa Cruz Biotechnology | Cat# sc-7947; RRID: AB_2121235 |
| Rabbit polyclonal anti-SEL1L | Homemade [1] | N/A |
| Rabbit polyclonal anti-HRD1/SYVN1 | Proteintech | Cat# 13473-1-AP; RRID: AB_2287023 |
| Rabbit monoclonal anti-ATG7 (D12B11) | Cell Signaling Technology | Cat# 8558S; RRID: AB_10831194 |
| Rabbit polyclonal anti-LC3 | Proteintech | Cat# 14600-1-AP; RRID: AB_2137737 |
| Rabbit polyclonal anti-BiP/GRP78 | Abcam | Cat# ab21685; RRID: AB_2119834 |
| Rabbit polyclonal anti-OS9 | Abcam | Cat# ab109510; RRID: AB_2864354 |
| Rabbit monoclonal anti-IRE1α (14C10) | Cell Signaling Technology | Cat# 3294; RRID: AB_823545 |
| Rabbit monoclonal anti-PERK (C33E10) | Cell Signaling Technology | Cat# 3192; RRID: AB_2095847 |
| Rabbit polyclonal anti-VAPA | Proteintech | Cat# 15275-1-AP; RRID: AB_2256991 |
| Rabbit polyclonal anti-VAPB | Proteintech | Cat# 14477-1-AP; RRID: AB_2288297 |
| Rabbit monoclonal anti-DRP1 (D6C7) | Cell Signaling Technology | Cat# 8570; RRID: AB_10950498 |
| Rabbit monoclonal anti-phospho-DRP1 (Ser616) | Cell Signaling Technology | Cat# 3455; RRID: AB_2085352 |
| Rabbit monoclonal anti-OPA1 (D6U6N) | Cell Signaling Technology | Cat# 80471; RRID: AB_2734117 |
| Rabbit polyclonal anti-MFN1 | Proteintech | Cat# 13798-1-AP; RRID: AB_2266318 |
| Rabbit monoclonal anti-MFN2 (D2D10) | Cell Signaling Technology | Cat# 9482; RRID: AB_2716838 |
| Mouse monoclonal anti-OXPHOS cocktail | Abcam | Cat# ab110413; RRID: AB_2629281 |
| Rabbit polyclonal anti-SigmaR1 | Proteintech | Cat# 15168-1-AP; RRID: AB_2301712 |
| Mouse monoclonal anti-α-tubulin (B-7) | Santa Cruz Biotechnology | Cat# sc-5286; RRID: AB_628411 |
| Guinea pig polyclonal anti-p62/SQSTM1 | MBL | Cat# PM066; RRID: AB_10952738 |
| Mouse monoclonal anti-ubiquitin (P4D1) | Santa Cruz Biotechnology | Cat# sc-8017; RRID: AB_628423 |
| Rabbit polyclonal anti-eIF2α | Cell Signaling Technology | Cat# 9722S; RRID: AB_2230924 |
| Rabbit polyclonal anti-phospho-eIF2α (Ser51) | Cell Signaling Technology | Cat# 9721S; RRID: AB_330951 |
| Rabbit polyclonal anti-TOMM20 | Proteintech | Cat# 11802-1-AP; RRID: AB_2207533 |
| Rabbit monoclonal anti-Perilipin1 (D1D8) | Cell Signaling Technology | Cat# 9349S; RRID: AB_2728690 |
| Goat Anti-Rabbit IgG (H+L), HRP-conjugated | Bio-Rad | Cat# 1706515; RRID: AB_11125142 |

| | | |
|---|---|---|
| Goat Anti-Mouse IgG (H+L), HRP-conjugated | Bio-Rad | Cat# 1706516; RRID: AB_11125547 |
| Goat Anti-Guinea Pig IgG (H+L), HRP-conjugated | Bio-Rad | Cat# AHP863P; RRID: AB_322395 |
| Goat Anti-Mouse IgG, 12 nm Colloidal Gold | Jackson ImmunoResearch | Cat# 715-205-150; RRID: AB_2340813 |
| Donkey Anti-Rabbit IgG, Alexa Fluor 488 | Jackson ImmunoResearch | Cat# 711-545-152; RRID: AB_2340846 |
| Donkey Anti-Guinea Pig IgG, Alexa Fluor 647 | Jackson ImmunoResearch | Cat# 706-605-148; RRID: AB_2340475 |
| Experimental models: Cell lines | | |
| WT, *Sel1L*$^{-/-}$, *Atg7*$^{-/-}$, *Sel1L*$^{-/-}$;*Atg7*$^{-/-}$ (DKO) brown preadipocytes | This study | N/A |
| HEK293T | ATCC | Cat# CRL-11268 |
| Experimental models: Organisms/strains | | |
| *Sel1L*$^{flox/flox}$ | Previous publication [2] | N/A |
| *Atg7*$^{flox/flox}$ | Previous publication [3] | N/A |
| *Ucp1-Cre* | The Jackson Laboratory | Cat# 024670 |
| *Sel1L*$^{flox/flox}$;*Ucp1-Cre (Sel1L*$^{Ucp1Cre}$*)* | Previous publication [1] | N/A |
| *Sel1L*$^{flox/flox}$;*Atg7*$^{flox/flox}$;*Adipo-Cre (Sel1;Atg7*$^{AdipoCre}$*)* | Previous publication [4] | |
| *Ire1a*$^{flox/flox}$ | Previous publication [5] | N/A |
| *Atg7*$^{flox/flox}$;*Ucp1-Cre (Atg7*$^{Ucp1Cre}$*)* | This study | N/A |
| *Sel1L*$^{flox/flox}$;*Atg7*$^{flox/flox}$;*Ucp1-Cre* (DKO) | This study | N/A |
| *Sel1L*$^{flox/flox}$;*Ire1a*$^{flox/flox}$;*Ucp1-Cre (Sel1L;Ire1α*$^{Ucp1Cre}$*)* | This study | N/A |
| Recombinant DNA | | |
| pCDNA3.1(+)-mito-GCaMP6f | Ming-Feng Tsai (UVA) | N/A |
| pHAGE2-pEF1α-mito-GCaMP6f | This study | N/A |
| pHAGE2-pEF1α-mito-DsRed | This study | N/A |
| pHAGE2-pEF1α-mito-mEmerald | This study | N/A |
| mEmerald-Sec61b-C1 | Addgene | Cat# 90992; RRID: Addgene_90992 |
| pLV-mito-DsRed | Addgene | Cat# 44386; RRID: Addgene_44386 |
| pRSV-Rev (REV) | Addgene | Cat# 12253; RRID: Addgene_12253 |
| pMDLg/pRRE (gag/pol) | Addgene | Cat# 12251; RRID: Addgene_12251 |
| pMD2.G (VSV-G) | Addgene | Cat# 12259; RRID: Addgene_12259 |
| Software and algorithms | | |
| Image Lab | Bio-Rad | https://www.bio-rad.com/en-us/product/image-lab-software?ID=KRE6P5E8Z |
| ImageJ | NIH | https://imagej.nih.gov/ij/ |
| GraphPad Prism 10 | GraphPad Software | RRID:SCR_002798 |
| Knossos | Open Source | https://knossos.app/ |
| Blender | Open Source | https://www.blender.org/ |
| Imaris 10.1 | Oxford Instruments | RRID: SCR_007370 |
| 3dEMtrace | Ariadne | https://ariadne.ai/ |
| Chemicals, Peptides, and Recombinant Proteins | | |
| 4-Hydroxytamoxifen | Cayman Chemical | Cat# 14854 |

| | | |
|---|---|---|
| Cycloheximide (CHX) | Cayman Chemical | Cat# 14126 |
| FCCP | Sigma-Aldrich | Cat# C2920 |
| Polyethylene glycol (PEG) 1,500 | ThermoFisher Scientific | Cat# A1624130 |
| Critical Commercial Assays | | |
| TRI reagent | Molecular Research Center | Cat# TR 118 |
| BCP phase separation reagent | Molecular Research Center | Cat# BP151 |
| Fluo-4/AM | ThermoFisher Scientific | Cat# F14201 |
| Pierce BCA Protein Assay Kit | ThermoFisher Scientific | Cat# 23227 |

### EXPERIMENTAL METHODS

#### Animals

All the mouse strains used in this study were on a C57BL/6J genetic background. $Sel1L^{flox/flox}$ [2], $Atg7^{flox/flox}$ [3], and brown adipocyte-specific $Sel1L^{Ucp1Cre}$ ($Sel1L^{flox/flox}$;*Ucp1-Cre*) [1], and $Ire1a^{flox/flox}$ [5] have been described previously. $Atg7^{Ucp1Cre}$ ($Atg7^{flox/flox}$;*Ucp1-Cre*) mice were generated by crossing $Atg7^{flox/flox}$ mice with *Ucp1-Cre* mice (The Jackson Laboratory, #024670). Double-knockout mice $Sel1L;Atg7^{Ucp1Cre}$ ($Sel1L^{flox/flox};Atg7^{flox/flox}$;*Ucp1-Cre*, DKO), $Sel1L;Atg7^{AdipCre}$ ($Sel1L^{flox/flox};Atg7^{flox/flox}$;*Adipoq-Cre*) [49], and $Sel1L;Ire1\alpha^{Ucp1Cre}$ ($Sel1L^{flox/flox};Ire1a^{flox/flox}$;*Ucp1-Cre*) were generated by intercrossing the single-knockout. $Sel1L^{flox/flox};Atg7^{flox/flox}$, $Sel1L^{flox/+};Atg7^{flox/flox}$, $Sel1L^{flox/flox};Atg7^{flox/+}$ and $Sel1L^{flox/+};Atg7^{flox/+}$ littermates were used as wildtype (WT) controls for DKO mice. $Sel1L^{flox/flox};Ire1\alpha^{flox/flox}$, $Sel1L^{flox/+};Ire1\alpha^{flox/flox}$, $Sel1L^{flox/flox};Ire1\alpha^{flox/+}$ and $Sel1L^{flox/+};Ire1\alpha^{flox/+}$ littermates were used as WT controls for $Sel1L;Ire1\alpha^{Ucp1Cre}$ mice. Experiments were performed using age- and sex-matched littermates between 10 and 12 weeks of age. All mice were housed in a specific pathogen-free animal facility at 22 ± 1 °C under a 12-hours light/dark cycle with 40-60% humidity and fed a low-fat diet (13% fat, 57% carbohydrate, 30% protein, LabDiet 5LOD), unless otherwise indicated. All animal procedures were approved by the Institutional Animal Care and Use Committee of the University of Michigan Medical School (PRO00010658), and University of Virginia (Protocol #4459), and in accordance with the National Institutes of Health (NIH) guidelines. Our study examined male and female animals, and similar findings were observed between sexes.

#### Animal experiments

Acute cold exposure was performed as previously described [1]. Mice were transferred from 22°C to 4°C, single-housed without food and nesting material, with ad libitum access to water. Core body temperature was measured at indicated time points using a mouse rectal probe (Braintree Scientific, Inc).

#### Western blot

Mouse tissues and cells were harvested, snap-frozen in liquid nitrogen and processed for Western blotting as previously described [1]. Primary antibody used included: anti-HSP90 (1:10,000; Santa Cruz, Cat# sc-7947), anti-SEL1L (1:10,000; homemade), anti-HRD1 (1:1000; Proteintech, Cat# 13473-1-AP), anti-ATG7 (1:1000; Cell Signaling, Cat# 8558S), anti-LC3 (1:1000; Proteintech, Cat# 14600-1-AP), anti-BiP (1:2000; Abcam, Cat# ab21685), anti-OS9 (1:2000; Abcam, Cat# ab109510), anti-IRE1α (1:1000; Cell Signaling, Cat# 3294), anti-PERK (1:1000; Cell Signaling, 3192), anti-VAPA (1:1000; Proteintech, Cat# 15275-1-AP), anti-VAPB (1:1000; Proteintech, Cat# 14477-1-AP), anti-DRP1 (1:1000; Cell Signaling, Cat# 8570), anti-p-DRP1 (Ser616) (1:1000; Cell Signaling, Cat# 3455), anti-OPA1 (1:1000; Cell Signaling, Cat# 80471), anti-MFN1 (1:1000; Proteintech, Cat# 13798-1-AP), anti-MFN2 (1:1000; Cell Signaling, Cat# 9482), anti-OXPHOS (1:1000; Abcam, Cat# ab110413), anti-Ubiquitin (1:500; Santa Cruz, Cat# sc-8017), anti-α-tubulin (1:500; Santa Cruz, Cat# sc2-5286), anti-p62 (1:1000; MBL, Cat# PM066), anti-eIF2α (1:1000; Cell Signaling, Cat# 9722S), anti-p-eIF2α (Ser51) (1:1000; Cell Signaling, Cat# 9721S), anti-Cleaved Caspase, anti-SigmaR1 (1:1000; Proteintech, Cat# 15168-1-AP). Membranes were washed with TBST and incubated with HRP-conjugated anti-Mouse (1:5000; Bio-Rad, Cat# 1706516), anti-Rabbit (1:5000; Bio-Rad, Cat# 1706515), or anti-Guinea Pig (1:5000; Bio-Rad, Cat# AHP863P) secondary antibodies at room temperature for 1 hour for ECL chemiluminescence detection system (Bio-Rad, Cat# 1705061) development. Band intensity was quantified using Image lab (Bio-Rad).

#### Plasmids

Mito-DsRed was amplified by PCR from pLV-mito-DsRed (Addgene; 44386), and mito-GCaMP6f was amplified by PCR from pCDNA3.1(+)-mito-GCaMP6f (a gift from Ming-Feng Tsai, University of Virginia). Both were cloned into the XbaI/BamHI sites of pHAGE2-pEF1α-eGFP (a gift from Yanzhuang Wang, Shenzhen Bay Laboratory, China) to generate the pHAGE2-pEF1α- mito-GCaMP6f and pHAGE2-pEF1α-mito-DsRed plasmids. Mito-mEmerald was amplified by overlapping PCR using pLV-mito-DsRed (Addgene, Cat# 44386) and mEmerald-Sec61b-C1

(Addgene, Cat# 90992) as templates and cloned into the XbaI/BamHI sites of pHAGE2-pEF1α-eGFP to generate the pHAGE2-pEF1α-mito-mEmerald plasmid.

#### Cell lines

WT and inducible *Sel1L*-deficient (*Sel1L*$^{-/-}$), *Atg7*$^{-/-}$, and *Sel1L*$^{-/-}$;*Atg7*$^{-/-}$ (DKO) brown pre-adipocytes were isolated from brown adipose tissue of mice and immortalized using SV40 large T antigen and differentiated into brown adipocytes as previously described [1,6]. All cells, including HEK293T cells were cultured at 37 °C with 5% $CO_2$ in DMEM with 10% fetal bovine serum (FBS) (Fisher Scientific, Cat# FB12999102). Cells were passaged every 2-3 days, when 80% confluence was reached. Cells were treated with 400 nM 4-hydroxytamoxifen (Cayman Chemical, Cat# 14854) starting on day three of differentiation for four consecutive days to generate knockout brown adipocytes.

#### PEG fusion assay

Lentiviral particles were generated by transfecting of HEK293T cells with Lipofectamine 3000 (Invitrogen, Cat# L3000008) and plasmids encoding pRRE (gag/pol) (Addgene, Cat# 12251), VSV-G (Addgene, Cat# 12259) and REV (Addgene, Cat# 12253), and either mito-mEmerald or mito-DsRed. Culture medium containing lentivirus were harvested at 24- and 48-hours post-transfection and used to infect brown preadipocytes. At 72 hours post-infection, preadipocytes stably expressing mito-mEmerald or mito-DsRed were sorted by flow cytometry. The PEG fusion assay was performed as described previously [7]. WT, *Sel1L*$^{-/-}$, *Atg7*$^{-/-}$, and DKO pre-adipocytes expressing either mito-mEmerald or mito-DsRed were cultured at 1:1 ratio, seeded in chambered coverslip µ-Slide 8 Well (Ibidi, Cat# 80826) and differentiated. Cells were treated with 4-hydroxytamoxifen for 4 consecutive days to induce gene deletion. On day 6 of differentiation, the cells were washed once with phosphate-buffered saline (PBS) and incubated in serum-free DMEM containing 33 µg/mL cycloheximide (CHX) (Cayman Chemical, Cat# 14126) for 30 min to inhibit protein synthesis. Cells were then treated with 50% (wt/vol) PEG 1,500 in serum-free DMEM for 2 min. Following polyethylene glycol (PEG) 1,500 (Thermo Fisher Scientific, Cat# A1624130) treatment, the cells were washed three times with DMEM supplemented with 10% FBS and CHX, then incubated for additional 4 hours in the same medium. Finally, cells were fixed in 4% paraformaldehyde for 20 min at room temperature, stained with DAPI, washed with PBS, and subjected to confocal imaging. Quantification of mitochondrial fusion was based on the colocalization of mito-DsRed (red) and mito-mEmerald (green) signals. Overlapping red and green signals resulting in yellow fluorescence were interpreted as evidence of mitochondrial content mixing in fused cells. Cell hybrids were classified into three categories: nearly complete overlap of red and green signals across the mitochondrial network (Full fusion); limited overlap of red and green signals (Partial fusion); spatially distinct red and green mitochondria without detectable colocalization (No fusion).

### Transmission electron microscopy (TEM)

TEM was performed as previously described[1]. Briefly, mice were anesthetized and perfused with 2.5% glutaraldehyde (Electron Microscopy Sciences, EMS, Cat# 16120), 4% paraformaldehyde (EMS, Cat# 15710) in 0.1 M sodium cacodylate buffer (EMS, Cat# 11653). BAT was immediately dissected and post-fixed overnight at 4 °C in the perfusion buffer. The fixed tissue was trimmed into 1-2 $mm^3$ cubes and then submitted to the University of Michigan Microscopy Core for further processing including washing, embedding and ultrathin sectioning (70 nm). The grid containing tissue sections was imaged using transmission electron microscope JEOL 1400 plus (Microscopy Core, University of Michigan Medical School) and FEI Tecnai F20 (Electron Microscopy Core, University of Virginia).

### Focused ion beam-SEM (FIB-SEM)

FIB-SEM was performed as previously described[1]. Briefly, fixed BAT cubes were submitted to the University of Michigan Microscopy Core for embedding. Embedded samples were analyzed at the Michigan Center for Materials Characterization (University of Michigan) using a FEI Helios Nanolab 650 DualBeam. The region of interest was determined, and the sample block was sequentially sectioned at 5 nm intervals. The resulting image stack was used for tomographic reconstruction. The ER and mitochondria were segmented using the 3dEMtrace platform of ariadne.ai (https://ariadne.ai/). briefly, Ground truth datasets were generated by manually annotating organelles in raw FIB-SEM images. Deep convolutional neural networks were trained on this manually annotated data and fine-tuned to achieve high-quality automated segmentation in 3D. The segmentations were proofread and refined by experts using the open-source software Knossos (https://knossos.app/). Meshes and binary TIFF masks for each segmentation class were generated for rendering and visualization. These outputs were analyzed using Knossos and further visualized using Blender, an open-source 3D visualization tool (https://www.blender.org/).

Lipid droplets, nucleus, and blood vessels segmentation was performed using Gradient Boosted Tree (GBT) pixel classifier. In brief, semantic segmentation of ER and instance segmentation of mitochondria were first converted to binary masks with the same dimension and voxel size as the raw data. Those masks were used to exclude ER and mitochondria regions from the subsequent pixel classifier training. The resulting masked 3D data were smoothed using a two-pixel gaussian filter followed by manual pixel-level annotation. GBT algorithm was implemented to train the pixel classifier for segmentation of lipid droplets, nucleus, and blood vessels, respectively. Iterative training and annotations were followed by manual examination. Final segmentation was analyzed and further visualized using Imaris software 10.1 (Oxford Instruments).

#### Immunogold labeling for TEM

Immunogold labeling was performed as previously described[1]. Briefly, BAT was fixed in buffer containing 3% formaldehyde and 0.2 M sucrose in 0.1 M Sorensen's phosphate buffer (pH 7.2, EMS, Cat# 11600-40), cut into small pieces, and processed by the University of Michigan Microscopy Core. Ultrathin sections (80 nm) were placed on bare nickel grids and stored at 4 °C. For immunogold labeling, grids were quenched in 80 mM glycine and incubated in blocking solution (EMS, Cat# 25599) containing 0.2% Tween-20 for 1 hour at room temperature followed by overnight incubation with primary antibody: anti-BiP (1:50; Abcam, Cat# ab21685) and anti-p62 (1:50, MBL, Cat# PM066) at 4 °C in a humidifying chamber. The grids were next incubated with colloidal gold-conjugated anti-Mouse secondary antibody (1:25; Jackson ImmunoResearch, Cat# 715-205-150) and anti-Guinea Pig (1:25; Jackson ImmunoResearch, Cat# 706-215-148) for 1 hour at room temperature. Post-fixation was performed with 1% glutaraldehyde and contrast staining was done using 0.5% uranyl acetate (EMS, Cat# 22400-2). After carbon coating, the grids were imaged using JEOL 1400 plus electron microscope (Microscopy Core, University of Michigan Medical School).

#### Histology and Immunofluorescence staining

Histology and immunofluorescence staining were performed as previously described [1]. Anesthetized mice were perfused with 10 ml of PBS, followed by 40 ml of 4% paraformaldehyde in PBS for fixation. BAT was dissected and post-fixed in 4% paraformaldehyde in PBS for 2 hours at 4°C. For hematoxylin and eosin (H&E) staining, samples were dehydrated, embedded in paraffin, sectioned, and stained at the Histology Core of University of Michigan Medical School and University of Virginia. For immunofluorescence staining of cryosections, samples were cryoprotected in 30% sucrose overnight, embedded in Tissue-Tek O.C.T. compound (Sakura, Cat# 4583) and frozen. Cryosections (7 μm thick) were prepared using a cryotome. Sections were washed with PBS, blocked in a solution of 10% normal donkey serum with 0.2% Triton X-100 in PBS for 1 hour at room temperature, and incubated overnight at 4°C with primary antibodies: anti-

Tomm20 (1:250; Proteintech, Cat# 11802-1-AP), anti-Perilipin (1:250; Cell Signaling Technology, Cat# 9349S), and anti-p62 (1:200, MBL, Cat# PM066). Then, sections were incubated in Alexa fluor 488 affinipure donkey anti-rabbit IgG (H+L) secondary antibody (1:500, Jackson ImmunoResearch, Cat# 711-545-152) and Alexa Fluor 647-AffiniPure donkey Anti-Guinea Pig IgG (H+L) (1:500; Jackson ImmunoResearch, Cat# 706-605-148) for 1 hour at room temperature. Images were acquired using Leica Stellaris 8 Resonant FALCON Confocal Microscope Stellaris 8 (Michigan Diabetes Research Center Microscopy & Image Analysis Core Microscopy & Image Analysis Core, University of Michigan Medical School) and Leica STELLARIS 8 tauSTED Super-Resolution Microscope (Advanced Microscopy Core, University of Virginia).

### RT-PCR and qPCR

Total RNA was extracted from brown adipose tissue using TRI reagent (Molecular Research Center, Cat# TR 118) in combination with BCP phase separation reagent (Molecular Research Center, Cat# BP151). *Xbp1* splicing was assessed by RT-PCR using specific primers (F: 5′-ACGAGGTTCCAGAGGTGGAG-3′; R: 5′-AAGAGGCAACAGTGTCAGAG-3′) as previously described [8]. The ratio of spliced to total *Xbp1* was quantified using Image Lab software (Bio-Rad). For SYBR Green-based qPCR, gene expression levels were normalized to *L32*. Primer sequences were as follows:

*L32* F: 5′-GAGCAACAAGAAAACCAAGCA-3′; R: 5′-TGCACACAAGCCATCTACTCA-3′.
*Vapa* F: 5′-CGGCCCAACAGTGGGATTAT-3′; R: 5′-GGGGTCCGTCTTGTTTGGAT-3′.
*Vapb* F: 5′-ACGGCATCTCACAGGTCTTG-3′; R: 5′-GGAAGCCACGTGGGTAAGAA-3′.
*Sigmar1* F: 5′-GACAGTATGCGGGGCTGGACCA-3′;
R: 5′-ACAACTGTCTCTCCTGGGTAGAAGA-3′.

### Mitochondrial DNA quantification

Total DNA was extracted from ~10–30 mg of brown adipose tissue using a lysis buffer containing proteinase K, followed by RNase A treatment and optional phenol–chloroform purification, as previously described [9]. DNA was precipitated with ammonium acetate and isopropanol, washed with 70% ethanol, and resuspended in TE buffer. Mitochondrial and nuclear DNA copy numbers were determined by SYBR Green-based qPCR. Mitochondrial DNA copy numbers were normalized to Nucleus gene *Hk2*. Primer sequences were as follows:

*Mt16s rRNA* F: 5′-CTAGAAACCCCGAAACCAAA-3′; R: 5′-CCAGCTATCACCAAGCTCGT-3′;
*Mtcox1* F: 5′-CTACTATTCGGAGCCTGAGC-3′; R: 5′-GCATGGGCAGTTACGATAAC-3′;
*Hk2* F 5′-GCCAGCCTCTCCTGATTTTAGTGT-3′; R: 5′-GGGAACACAAAAGACCTCTTCTGG-3′.

### Measurement of cytoplasmic and mitochondrial Ca2+ signals

$Ca^{2+}$ measurement was performed as previously described [10]. Briefly, preadipocytes were seeded on 35 mm glass-bottom dishes and differentiated into adipocytes. Cytoplasmic and mitochondrial $Ca^{2+}$ signals were monitored by Leica Thunder imager. All recordings were performed in modified Tyrode's solution (TS) containing 130 mM NaCl, 5.4 mM KCl, 1 mM $MgCl_2$, 100 µM EGTA, 20 mM HEPES, and 5 mM glucose (pH 7.4, adjusted with NaOH). Cytoplasmic $Ca^{2+}$ signals were assessed using the cytosolic $Ca^{2+}$-sensitive probe Fluo-4/AM (maximum excitation/emission: 494/506 nm). Differentiated adipocytes grown on glass-bottom dishes were incubated with 5 µM Fluo-4/AM in complete medium for 60 minutes.

Mitochondrial $Ca^{2+}$ uptake following $Ca^{2+}$ release was assessed using mito-GCaMP6f, a mitochondria-targeted, genetically encoded $Ca^{2+}$ indicator (Kd = 375 nM; maximum excitation/emission: 488/500–550 nm). Preadipocytes stably expressing mito-GCaMP6f were generated via lentiviral transduction. Cells were washed twice with Tyrode's solution, and the baseline fluorescence was recorded prior to stimulation with 500 µM ATP to induce $Ca^{2+}$ release. Time-lapse imaging was performed by acquiring 20-30 frames at 5-second intervals using FITC filter (EX: 460-500; DC: 505; EM: 512-542) and a 40 × objective. Images were analyzed using ImageJ (NIH) and fluorescence intensity was quantified.

### Oxygen consumption rate (OCR)

Mitochondrial isolation was performed as previously described [1]. briefly, BAT was dissected, rinsed, and homogenized in 30 mL ice-cold homogenization buffer (HB) (70 mM sucrose, 210 mM mannitol, 5 mM MOPS, 5 mM KH2PO4, 1 mM EGTA and 0.5% (w/v) fatty acid-free BSA, pH 7.2) using Potter-Elvehjem tissue grinder. The homogenate was centrifuged at 8,000 *x* g, and the pellet was resuspended in HB without BSA. The supernatant was centrifuged at 800 g for 10 minutes at 4 °C to remove nuclei and debris. The resulting supernatant was centrifuged again at 8,000 g for 10 minutes at 4 °C to isolate the crude mitochondrial fraction, which was resuspended in a small volume of HB without BSA for protein concentration determination and OCR measurements. For OCR measurements, isolated mitochondria were resuspended in MiR05 buffer (Oroboros, Cat# 60101-01). OCR was measured using the OROBOROS Oxygraph-2K system. Basal respiratory rate was assessed in the presence of 5 mM pyruvate and 1 mM malate, and the maximal respiratory rate was determined by titrating carbonyl cyanide p-trifluoromethoxy phenylhydrazone (FCCP, Sigma-Aldrich, Cat# C2920).

### Blue-native PAGE

Blue native PAGE (BN-PAGE) was performed as previously described [9]. Briefly, mitochondria isolated from BAT were homogenized in solubilization buffer (1% digitonin, 50 mM NaCl, 50 mM imidazole, 1 mM EDTA, 2 mM 6-aminocaproic acid and protease inhibitors, pH 7.0). After centrifugation at 20,000 *x* g for 20 min at 4°C, Coomassie Brilliant Blue (final concentration: 0.2%) and glycerol (final concentration: 10%) were added to the supernatant. Samples were then subjected to electrophoresed on 4-13% polyacrylamide gradient gels, transferred to PVDF membranes, and analyzed by immunoblotting.

### Mitochondria-associated membranes (MAMs) isolation

MAMs were isolated from BAT as previously described [11]. Briefly, BAT was homogenized in homogenization buffer (225 mM mannitol, 75 mM sucrose, 0.5% BSA, 0.5 mM EGTA, and 30 mM Tris-HCl, pH 7.4). Homogenates were centrifuged at 750 *x* g for 5 min to remove unbroken cells and nucleus. The crude mitochondrial pellets were collected after centrifugation at 9, 000 *x* g for 10 min, following by washing two times by washing buffer (225 mM mannitol, 75 mM sucrose, 0.5% BSA and 30 mM Tris-HCl, pH 7.4) at 10, 000 *x* g. Then resulting crude mitochondria was resuspended in mitochondria resuspending buffer (MRB) (250 mM mannitol, 5 mM HEPES and 0.5 mM EGTA, pH 7.4), and were subjected to 30% Percoll gradient centrifugation at 9,500 *x* g for 30 min using a Beckman Coulter Optima L-100 XP Ultracentrifuge (SW40 rotor, Beckman) at 4°C. A low-density band (representing the MAM fraction) was resuspended in MRB and further purified by centrifugation at 6,300 *x* g for 10 min to removing contaminating pellets. The mitochondrial fraction in the high-density band was resuspended in MRB and pelleted by centrifugation at 6,300 *x* g for 10 min. The MAM fraction was obtained by ultracentrifugation at 100,000 *x* g for 1 hour at 4°C using a 70-Ti rotor (Beckman). Microsomes were pelleted from the mitochondrial supernatant by ultracentrifugation at 100,000 *x* g for 1 hour, and the cytosol fraction was collecting as the resulting supernatant. SDS-PAGE sample buffer was added to each fraction for analysis by Western blotting.

### Statistical analyses

All results are presented as mean ± standard error of the mean (SEM). Each experiment was independently repeated two to three times and/or included multiple independent biological replicates, from which representative data are shown. Statistical comparisons between two groups were performed using an unpaired two-tailed Student's *t*-test. For comparisons involving more than two groups, one-way or two-way analysis of variance (ANOVA) followed by Tukey's multiple comparisons test was applied. All statistical analyses were performed using GraphPad Prism 10 (GraphPad Software Inc.).

### REFERENCES


1. Zhou, Z., Torres, M., Sha, H., Halbrook, C.J., Van den Bergh, F., Reinert, R.B., Yamada, T., Wang, S., Luo, Y., Hunter, A.H., et al. (2020). Endoplasmic reticulum-associated degradation regulates mitochondrial dynamics in brown adipocytes. Science *368*, 54-60. 10.1126/science.aay2494.
2. Sun, S., Shi, G., Han, X., Francisco, A.B., Ji, Y., Mendonca, N., Liu, X., Locasale, J.W., Simpson, K.W., Duhamel, G.E., et al. (2014). Sel1L is indispensable for mammalian endoplasmic reticulum-associated degradation, endoplasmic reticulum homeostasis, and survival. Proc Natl Acad Sci U S A *111*, E582-591. 10.1073/pnas.1318114111.
3. Komatsu, M., Waguri, S., Ueno, T., Iwata, J., Murata, S., Tanida, I., Ezaki, J., Mizushima, N., Ohsumi, Y., Uchiyama, Y., et al. (2005). Impairment of starvation-induced and constitutive autophagy in Atg7-deficient mice. J Cell Biol *169*, 425-434. 10.1083/jcb.200412022.
4. Wu, S.A., Shen, C., Wei, X., Zhang, X., Wang, S., Chen, X., Torres, M., Lu, Y., Lin, L.L., Wang, H.H., et al. (2023). The mechanisms to dispose of misfolded proteins in the endoplasmic reticulum of adipocytes. Nature communications *14*, 3132. 10.1038/s41467-023-38690-4.
5. Iwawaki, T., Akai, R., Yamanaka, S., and Kohno, K. (2009). Function of IRE1 alpha in the placenta is essential for placental development and embryonic viability. Proc Natl Acad Sci U S A *106*, 16657-16662. 10.1073/pnas.0903775106.
6. Klein, J., Fasshauer, M., Klein, H.H., Benito, M., and Kahn, C.R. (2002). Novel adipocyte lines from brown fat: a model system for the study of differentiation, energy metabolism, and insulin action. Bioessays *24*, 382-388. 10.1002/bies.10058.
7. Nascimento, A., Lannigan, J., and Kashatus, D. (2016). High-throughput detection and quantification of mitochondrial fusion through imaging flow cytometry. Cytometry A *89*, 708-719. 10.1002/cyto.a.22891.
8. Sha, H., He, Y., Chen, H., Wang, C., Zenno, A., Shi, H., Yang, X., Zhang, X., and Qi, L. (2009). The IRE1alpha-XBP1 pathway of the unfolded protein response is required for adipogenesis. Cell Metab *9*, 556-564. 10.1016/j.cmet.2009.04.009.
9. Wittig, I., Braun, H.P., and Schagger, H. (2006). Blue native PAGE. Nat Protoc *1*, 418-428. 10.1038/nprot.2006.62.
10. Tsai, C.W., Rodriguez, M.X., Van Keuren, A.M., Phillips, C.B., Shushunov, H.M., Lee, J.E., Garcia, A.M., Ambardekar, A.V., Cleveland, J.C., Jr., Reisz, J.A., et al. (2022). Mechanisms and significance of tissue-specific MICU regulation of the mitochondrial calcium uniporter complex. Mol Cell *82*, 3661-3676 e3668. 10.1016/j.molcel.2022.09.006.
11. Wieckowski, M.R., Giorgi, C., Lebiedzinska, M., Duszynski, J., and Pinton, P. (2009). Isolation of mitochondria-associated membranes and mitochondria from animal tissues and cells. Nat Protoc *4*, 1582-1590. 10.1038/nprot.2009.151.

### FIGURES AND FIGURE LEGENDS

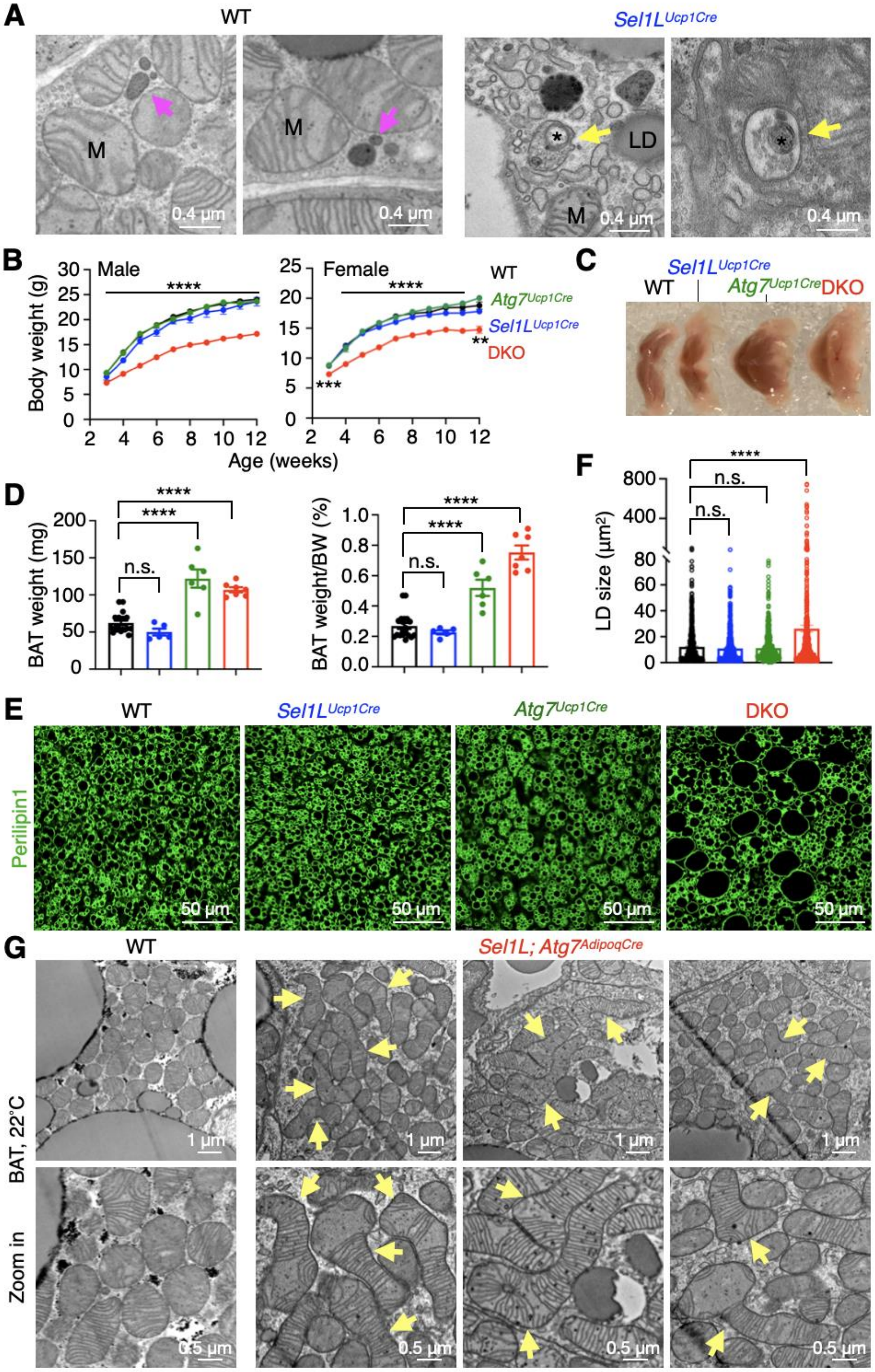

**Figure S1. Generation and characterization of DKO mice lacking SEL1L and ATG7 specifically in brown adipocytes, related to Figure 1.**

(A) Representative TEM images of BAT showing lysosomes (magenta arrows) and mitochondria-like structures (*) in the autophagosome (yellow arrows) from 12-week-old male littermates. LD, lipid droplet; M, mitochondrion.

(B) Growth curves of male (left) and female (right) littermates at different ages. Statistical comparison is between WT and DKO mice. n = 10-21 mice per time point, Two-way ANOVA.

(C) Representative pictures of BAT from 12-week-old male littermates.

(D) Quantification of BAT weight (left) and BAT weight normalized to body weight (BW) (right) from 12-week-old male littermates. n = 19 mice for WT, 5 for $Sel1L^{Ucp1Cre}$, 6 for $Atg7^{Ucp1Cre}$ and 7 for DKO, One-way ANOVA.

(E-F) Representative Perilipin1 staining of BAT from 12-week-old male littermates, with quantification of LDs size shown in (F). n = 783 LDs for WT, 704 for $Sel1L^{Ucp1Cre}$, 621 for $Atg7^{Ucp1Cre}$, and 769 for DKO, two mice per group, One-way ANOVA.

(G) Representative TEM images of BAT from 12-week-old male WT and DKO (*Sel1L*; $Atg7^{AdipoqCre}$) littermates housed at RT. Arrows point to megamitochondria. n = 2 mice per group.

Data are mean ± SEM. n.s., not significant; ****, $p < 0.0001$.

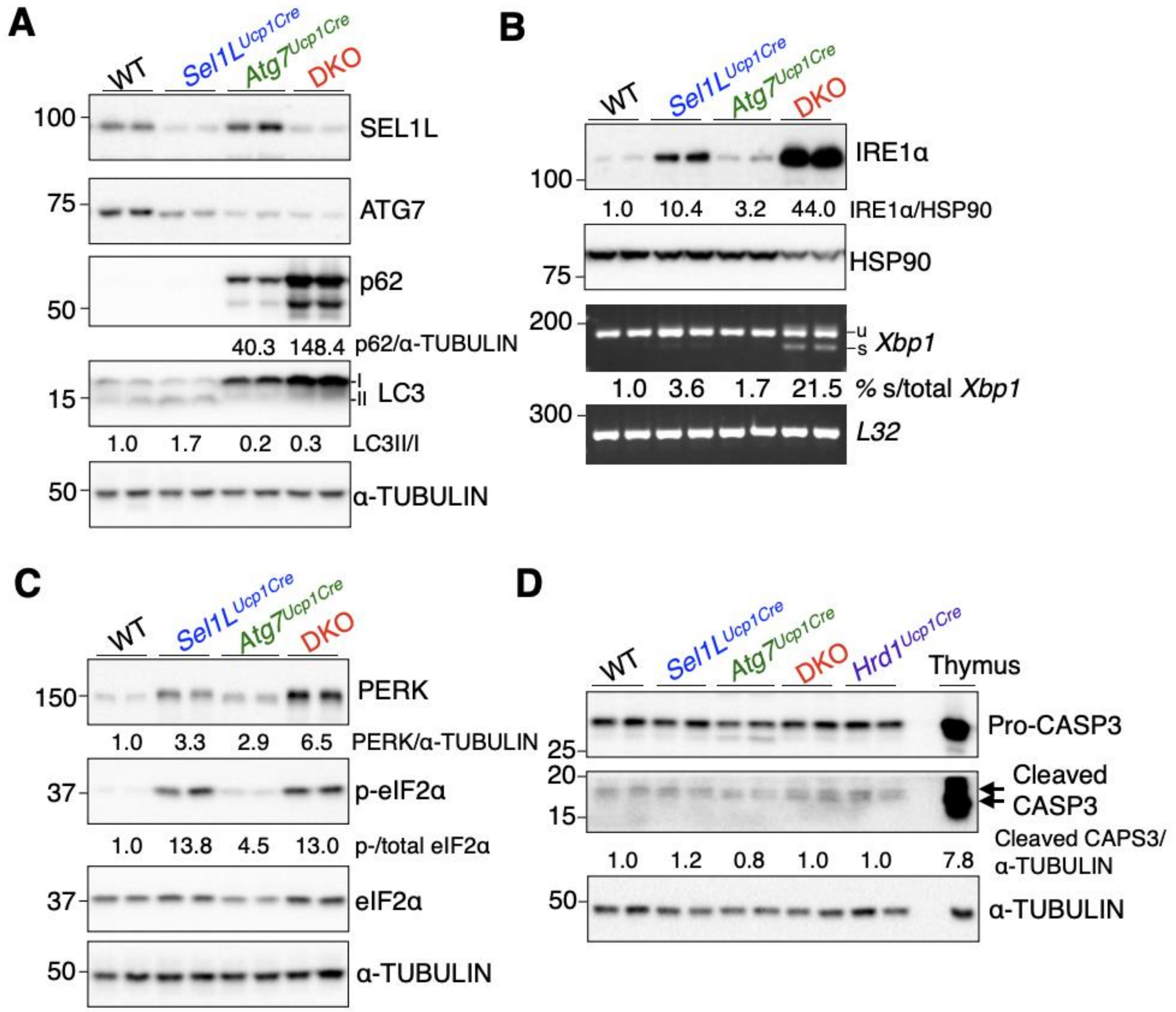


**Figure S2. Combined loss of SEL1L-HRD1 ERAD and autophagy in brown adipocytes leads to mild ER stress without inducing cell death, related to Figure 1.**

(A-C) Representative assays for autophagy (ATG7, p62, LC3) (A) and UPR markers (IRE1α, *Xbp1*, PERK, and p-eIF2α) (B and C) in BAT from 12-week-old littermates, with quantification shown below the blots.

(D) Immunoblot analysis of cleaved caspase-3 in BAT from 12-week-old littermates, with quantification shown below the blots. Mouse thymus was used as a positive control.

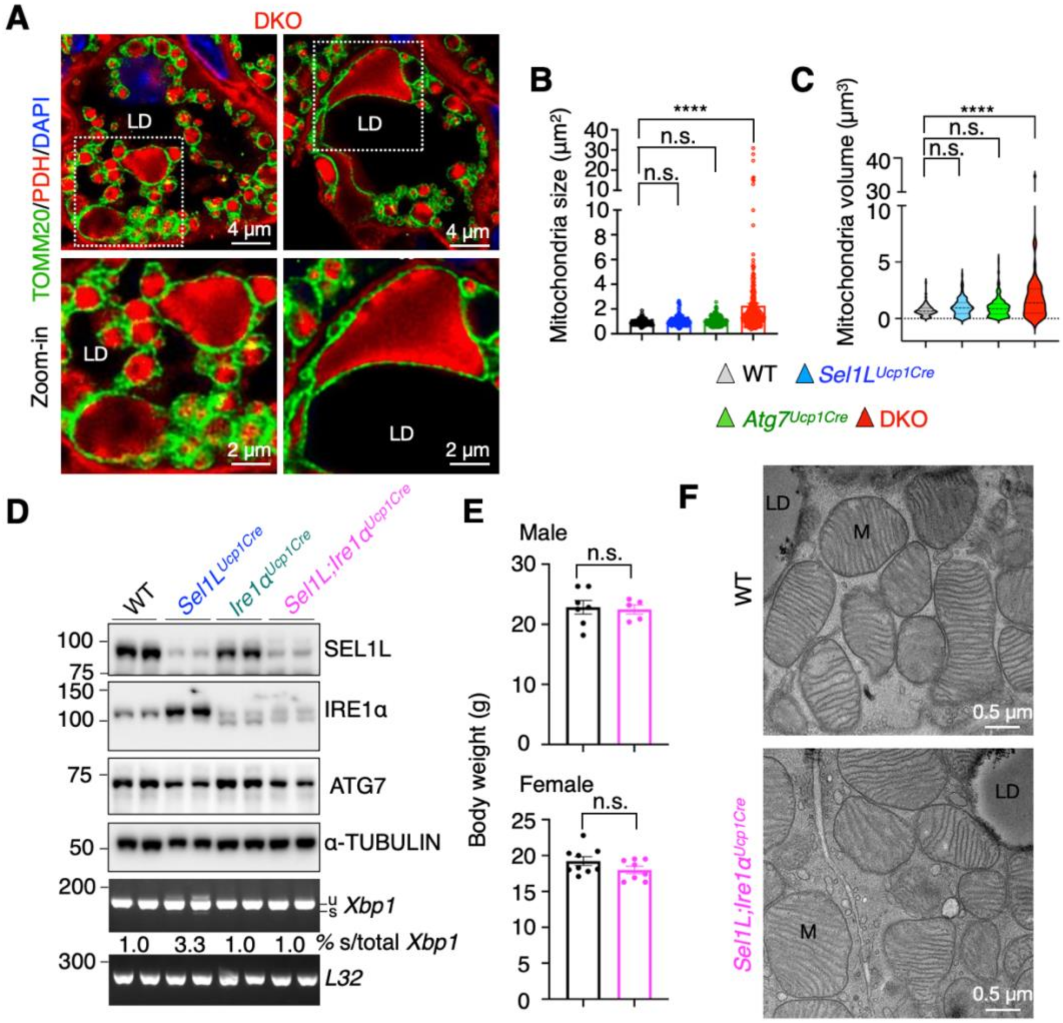


**Figure S3. Formation of megamitochondria in *Sel1L;Atg7*-deficient brown adipocytes, but not in *Sel1L;Ire1a*-deficient brown adipocytes, related to Figure 2.**

(A) Representative confocal immunofluorescent images of TOMM20 (mitochondrial outer membrane protein) and pyruvate dehydrogenase (PDH, mitochondrial matrix protein) staining in BAT sections from 12-week-old DKO male mice. Arrows point to megamitochondria. LD, lipid droplet. n=3 mice each group.

(B) Quantitation of mitochondrial size circled by TOMM20 in Figure 2A and S3A. n = 194 mitochondria for WT, 231 for *Sel1L*$^{Ucp1Cre}$, 165 for *Atg7*$^{Ucp1Cre}$, and 297 for DKO, One-way ANOVA.

(C) Quantification of mitochondrial volume reconstructed from FIB-SEM images shown in Figure 2B. n = 195 mitochondria for WT, 129 for *Sel1L*$^{Ucp1Cre}$, 283 for *Atg7*$^{Ucp1Cre}$, and 51 for DKO, One-way ANOVA.

(D) Immunoblot analysis of SEL1L, IRE1α, and ATG7 in BAT from 10-week-old littermates.

(E) Body weight of 10-week-old male (top) and female (bottom) littermates. n = 5-9 mice per genotype, Student's *t*-test.

(F) Representative TEM images of BAT from 10-week-old male littermates. LD, lipid droplet; M, mitochondrion. n = 2 mice per group.

Data are mean ± SEM. n.s., not significant; ****, $p < 0.0001$.

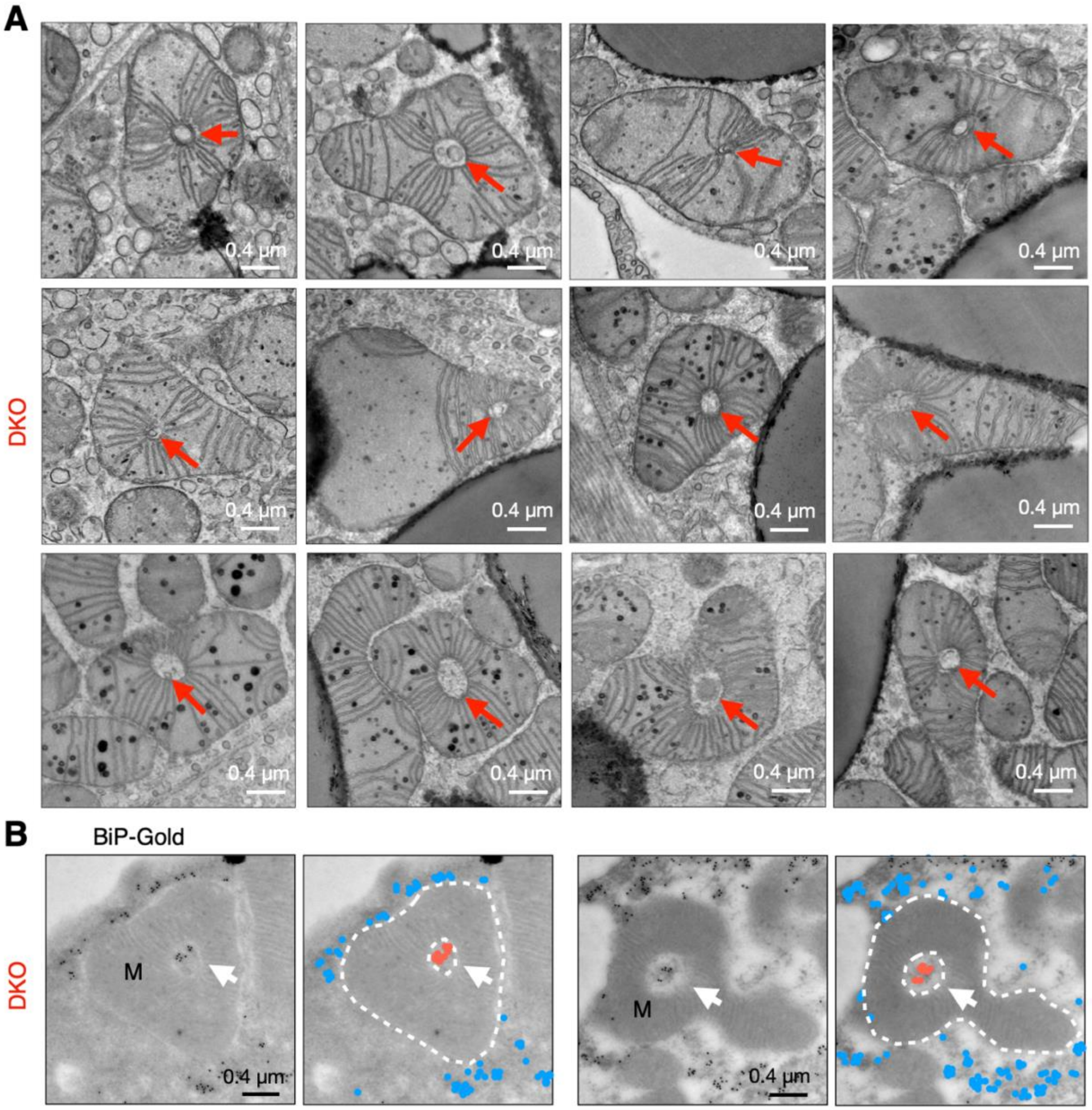


**Figure S4. ERAD and autophagy deficiency leads to the formation of megamitochondria with perforating ER tubules under RT, related to Figure 3.**

(A) Representative TEM images of BAT showing megamitochondria wrapping around the tubular structures from 12-week-old DKO mice. Arrows point to mitochondrion-perforating tubules. n = 5 mice.

(B) Representative TEM images of BAT following BiP-specific immunogold labelling from 12-week-old DKO mice, with pseudo colored blue and red dots shown on the right. White dotted lines outline megamitochondria. Arrows point to mitochondrion-perforating ER tubules. M, mitochondrion; n = 2 mice.

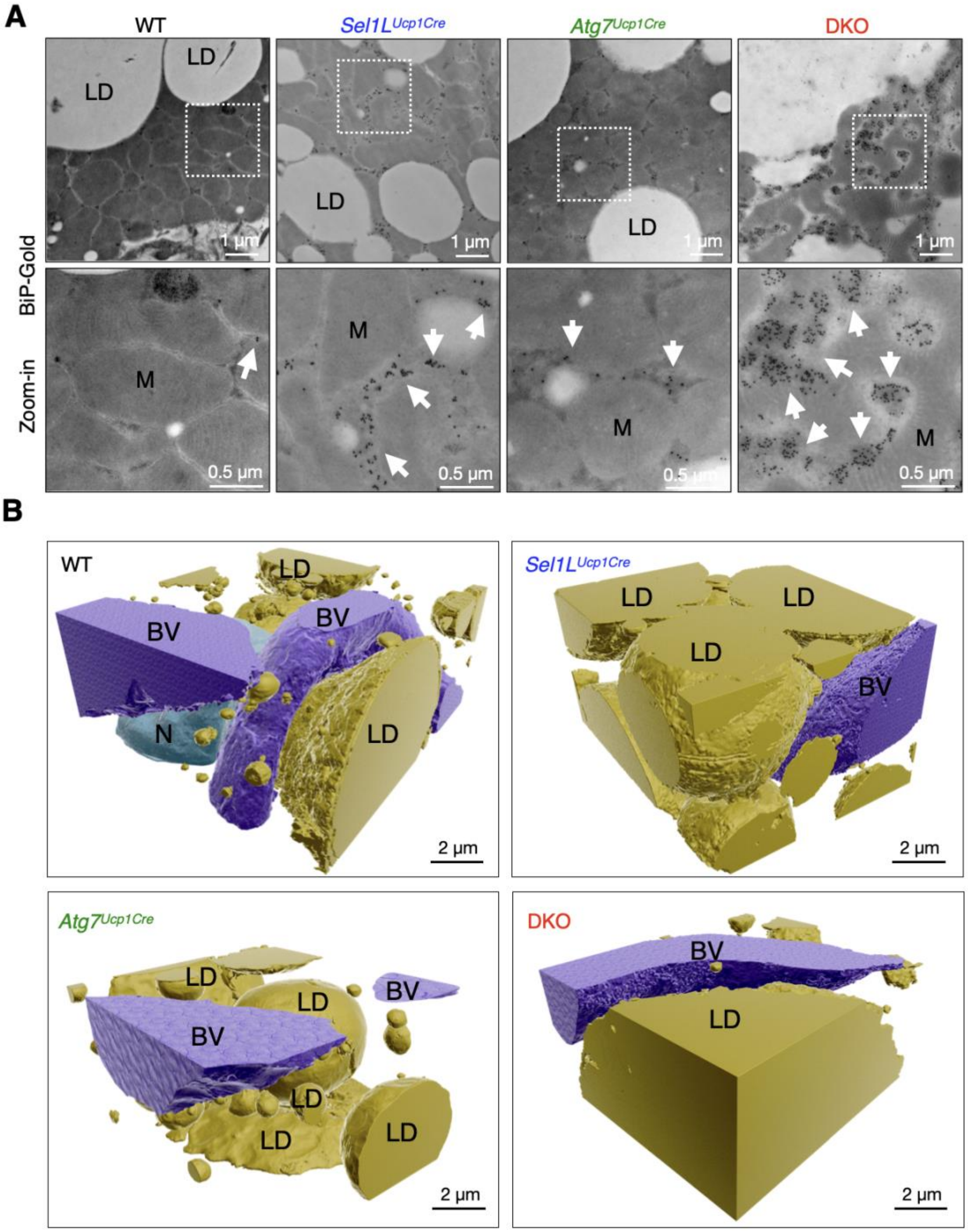


**Figure S5. ERAD and autophagy synergistically control the ER architecture in brown adipocytes, related to Figure 4.**

(A) Representative TEM images of BAT following BiP-specific immunogold labelling from 12-week-old male littermates . Arrows point to BiP-immunogold particles. LD, lipid droplet; M, mitochondrion. n = 2 mice per group.

(B) 3D reconstruction of FIB-SEM images of BAT showing lipid droplets (yellow), nucleus (green), and blood vessels (purple) from 12-week-old male littermates. LD, lipid droplet; N, Nucleus; BV, blood vessels.

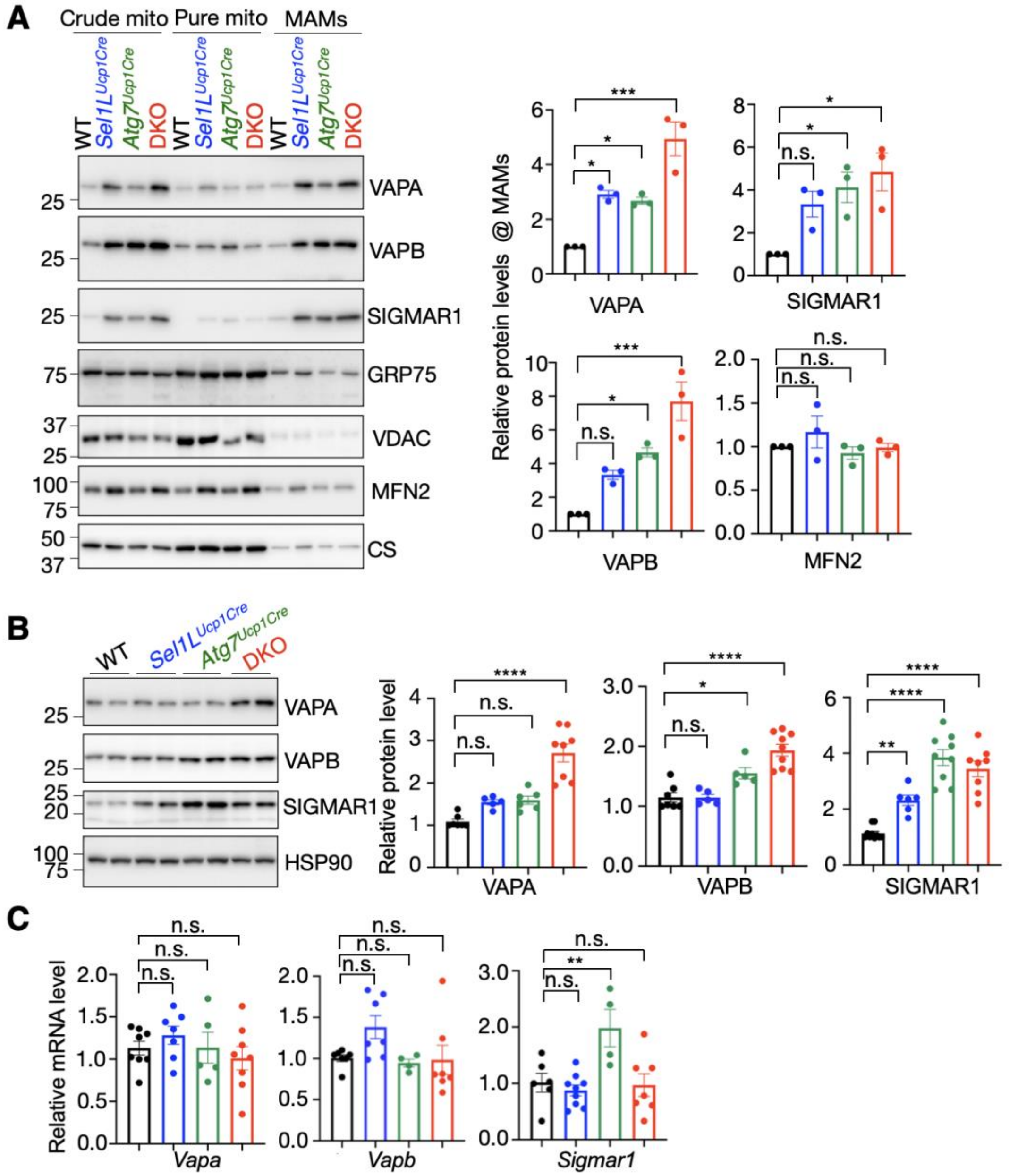


**Figure S6. ERAD and autophagy deficiency increases the abundance of a subset of MAMs-tethering proteins, related to Figure 5.**

(A) Immunoblot analysis of indicated proteins from subcellular fractionation in BAT from 12-week-old littermates, with quantification of indicated proteins (VAPA, VAPB, SIGMAR1, and MFN2 in MAMs shown right. CM: crude mitochondria, PM: pure mitochondria, MAM: Mitochondria associated membranes. n = 3 per group, One-way ANOVA.

(B) Immunoblot analysis of VAPA, VAPB, and SIGMAR1 in BAT from 12-week-old littermates, with quantification of proteins abundance normalized to HSP90 shown right. n = 5-9 per group, One-way ANOVA.

(C) qPCR analyses of *Vapa, Vapb, and Sigmar1* mRNA level in BAT. n = 4-9 per group, One-way ANOVA.

Data are mean ± SEM. n.s., not significant; **p* < 0.1; ***p* < 0.01; ****p* < 0.001; ****, *p* < 0.0001.

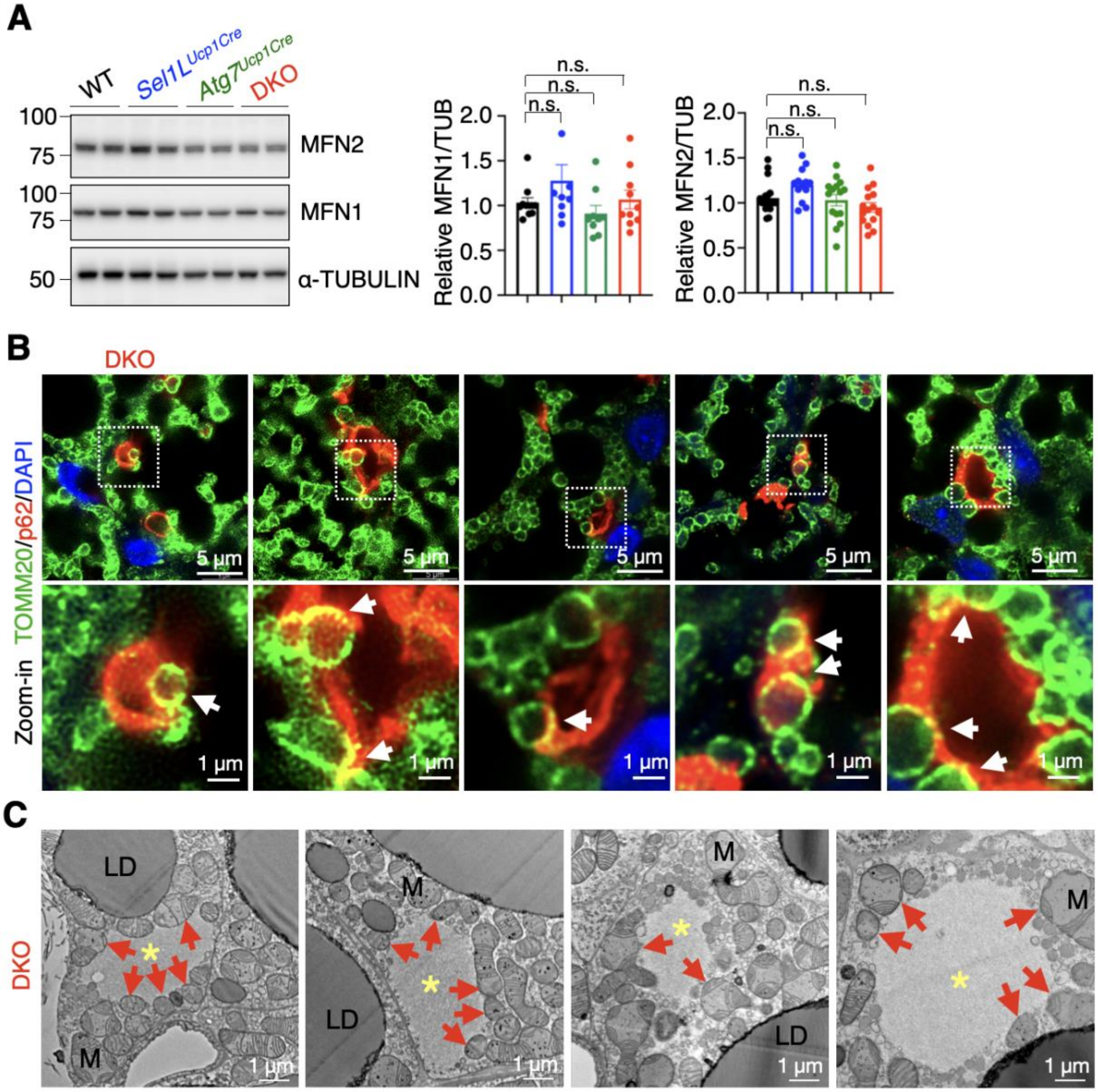


**Figure S7. ERAD and autophagy synergistically regulate p62 aggregation and mitochondrial quality control, related to Figure 6.**

(A) Immunoblot analysis of MFN1/2 as mitochondrial-fusion markers in BAT 12-week-old littermates, with quantification of MFN1/2 normalized to α-TUBULIN (TUB) shown on the right. n = 9-28 per genotype, One-way ANOVA.

(B) Representative immunofluorescent images showing labeling of Tomm20 (green) and p62 (red) in BAT sections from 12-week-old DKO male mice. White arrows mark the colocalization of p62 and Tomm20. n = 3 mice.

(C) Representative TEM images of p62 inclusions in BAT from 12-week-old DKO mice. Asterisks indicate p62 inclusion. Red arrows point to mitochondrion near p62 inclusion. LD, lipid droplet; M, mitochondrion. n = 5 mice.

Data are mean ± SEM. n.s., not significant.

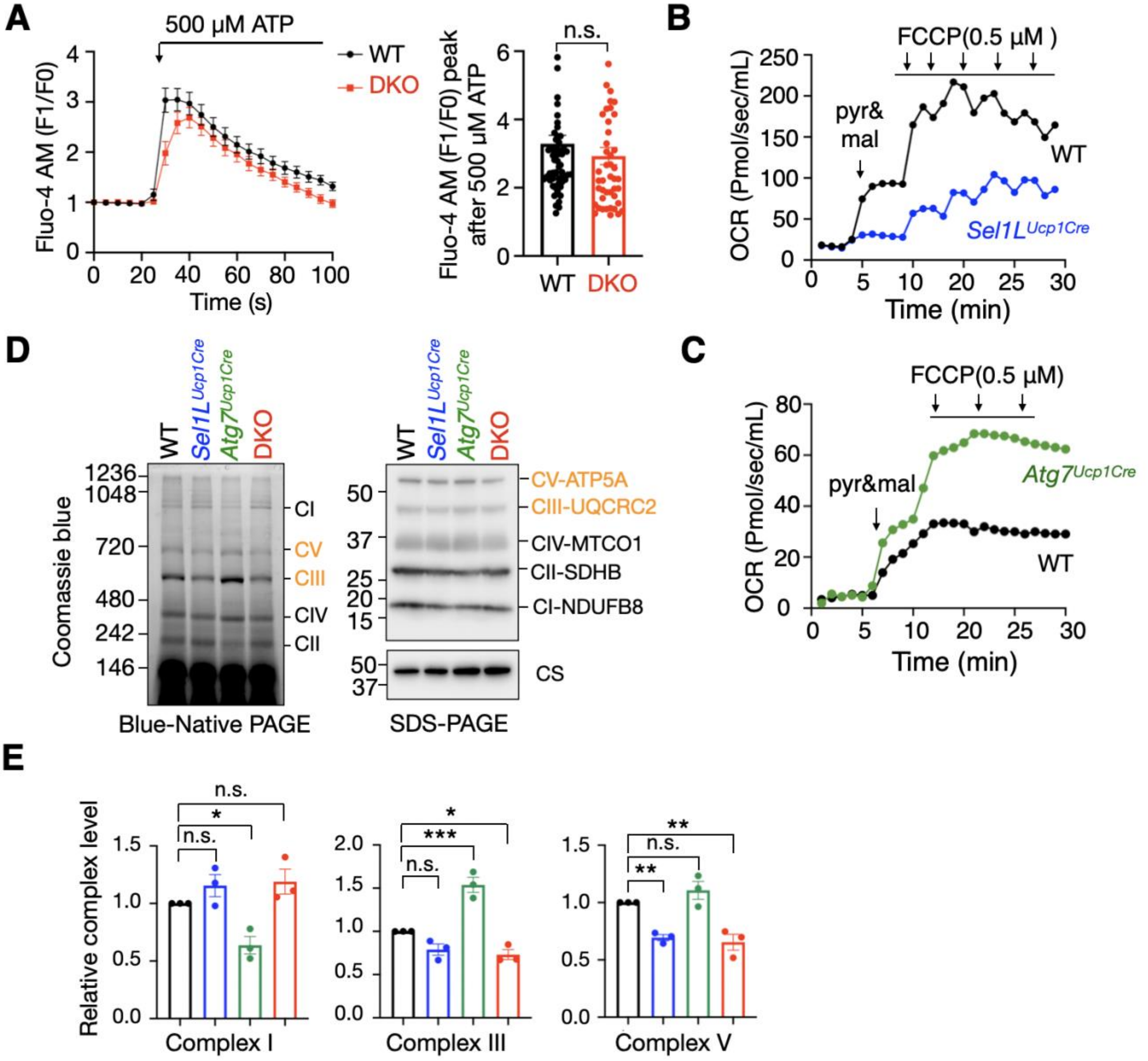


**Figure S8.** ***Sel1L;Atg7*** **deficiency compromises mitochondria function and thermogenesis in brown adipocytes, related to Figure 7.**

(A) Representative traces of cytosolic $Ca^{2+}$ responds in WT and DKO differentiated adipocytes challenged with 500 μM ATP using Fluo-4 AM. Quantification of peak Fluo-4 AM is shown on the right. n=62 cells for WT and 46 for DKO, three repeats, Student's *t*-test.

(B-C) Oxygen consumption rate (OCR) of purified mitochondria from BAT of 12-week-old littermates. Three repeats.

(D) Mitochondrial OXPHOS complex analysis of isolated mitochondria from BAT of 12-week-old littermates by Blue-native (BN) PAGE followed by Coomassie Blue staining (left panel) and SDS-PAGE following by immunoblotting (right panel). Three repeats.

(E) Quantification of complex I, complex III, and complex V from BN-PAGE normalized to CS from SDS-PAGE is shown on the below. n=3 per group, One-way ANOVA.

Data are mean ± SEM. n.s., not significant; **p* < 0.1; ***p* < 0.01; ****p* < 0.001.

### VIDEO LEGENDS

**Video S1–S4. Raw FIB-SEM volumes and 3D reconstructions of mitochondria, ER, and mitochondria-associated membranes (MAMs) in brown adipose tissue (BAT).**

Raw FIB-SEM image stacks and corresponding 3D renderings of mitochondria, endoplasmic reticulum (ER), and MAMs in BAT from 12-week-old male littermates. The datasets illustrate differences in mitochondrial architecture and ER–mitochondria interactions across genotypes, WT (video S1), $Sel1L^{Ucp1Cre}$ (video S2), $Atg7^{Ucp1Cre}$ (video S3), and DKO (video S4). color segmentations are indicated within each video.

### Table S1. Full List of Oligonucleotides Used in This Study

| Primer Name and Sequence | Application / Note |
|---|---|
| *Xbp1* F: ACGAGGTTCCAGAGGTGGAG | RT-PCR |
| *Xbp1* R: AAGAGGCAACAGTGTCAGAG | RT-PCR |
| *L32* F: GAGCAACAAGAAAACCAAGCA | RT-PCR and qPCR |
| *L32* R: TGCACACAAGCCATCTACTCA | RT-PCR and qPCR |
| *Vapa* F: CGGCCCAACAGTGGGATTAT | qPCR |
| *Vapa* R: GGGGTCCGTCTTGTTTGGAT | qPCR |
| *Vapb* F: ACGGCATCTCACAGGTCTTG | qPCR |
| *Vapb* R: GGAAGCCACGTGGGTAAGAA | qPCR |
| *Sigmar1* F: GACAGTATGCGGGGCTGGACCA | qPCR |
| *Sigmar1* R: ACAACTGTCTCTCCTGGGTAGAAGA | qPCR |
| *Mt16s rRNA* F: CTAGAAACCCCGAAACCAAA | qPCR |
| *Mt16s rRNA* R: CCAGCTATCACCAAGCTCGT | qPCR |
| *Mtcox1* F: CTACTATTCGGAGCCTGAGC | qPCR |
| *Mtcox1* R: GCATGGGCAGTTACGATAAC | qPCR |
| *Hk2* F: GCCAGCCTCTCCTGATTTTAGTGT | qPCR |
| *Hk2* R: GGGAACACAAAAGACCTCTTCTGG | qPCR |
